# Macromolecular crowding links ribosomal protein gene dosage to growth rate in *Vibrio cholerae*

**DOI:** 10.1101/619304

**Authors:** Alfonso Soler-Bistué, Sebastián Aguilar-Pierlé, Marc Garcia-Garcerá, Marie-Eve Val, Odile Sismeiro, Hugo Varet, Rodrigo Sieira, Evelyne Krin, Ole Skovgaard, Diego J. Comerci, Eduardo P. C. Rocha, Didier Mazel

## Abstract

Ribosomal protein (RP) genes locate near the replication origin (*oriC*) in fast-growing bacteria, which is thought to have been selected as a translation optimization strategy. Relocation of *S10-spc-α* locus (S10), which codes for most of the RP, to ectopic genomic positions shows that its relative distance to the *oriC* correlates to a reduction on its dosage, its expression, and bacterial growth rate. Deep-sequencing revealed that S10 relocation altered chromosomal replication dynamics and genome-wide transcription. Such changes increased as a function of *oriC*-S10 distance. Strikingly, in this work we observed that protein production capacity was independent of S10 position. Since RP constitute a large proportion of cell mass, lower S10 dosage could lead to changes in macromolecular crowding, impacting cell physiology. Accordingly, cytoplasm fluidity was higher in mutants where S10 is most distant from *oriC*. In hyperosmotic conditions, when crowding differences are minimized, the growth rate and replication dynamics were highly alleviated in these strains. Therefore, on top of its essential function in translation, RP genomic location contributes to sustain optimal macromolecular crowding. This is a novel mechanism coordinating DNA replication with bacterial growth.

## Introduction

Replication, gene expression and segregation are tightly coordinated with the cell cycle to preserve homeostasis (1, 2). Genome structure is a plausible factor contributing to integrate these many simultaneous processes occurring on the same template. The relative simplicity and the increasing amount of available data render bacterial genomes ideal models to study this subject (3–6).

Bacterial chromosomes are highly variable in their gene content, but highly conserved in terms of the order of core genes in the chromosomes. Replication begins at a sole replication origin (*oriC*), proceeding bidirectionally along two equally sized replichores until the terminal region (*ter*). This organizes the genome along an *ori-ter* axis that interplays with cell physiology (Fig. 1a) (4, 5, 7). For instance, essential genes are overrepresented in the replicative leading strand to avoid head-on collisions between the replication and transcription machineries (8). Large inversions occur preferentially symmetrically with respect to the *ori-ter* axis to avoid the emergence of replichore size imbalance (9, 10). Recent studies indicate that gene order within the chromosome may play a relevant role in harmonizing the genome structure with cell physiology. Remarkably, key genes coding for nucleoid associated proteins, RNA polymerase modulators, topoisomerases and energy production are arranged along the *ori-ter* axis following the temporal order of their expression during growth phases (11, 12). In addition, recent studies have showcased an increasing number of traits whose expression is influenced by the genomic position of its encoding genes (13–15).

**Figure 1:**
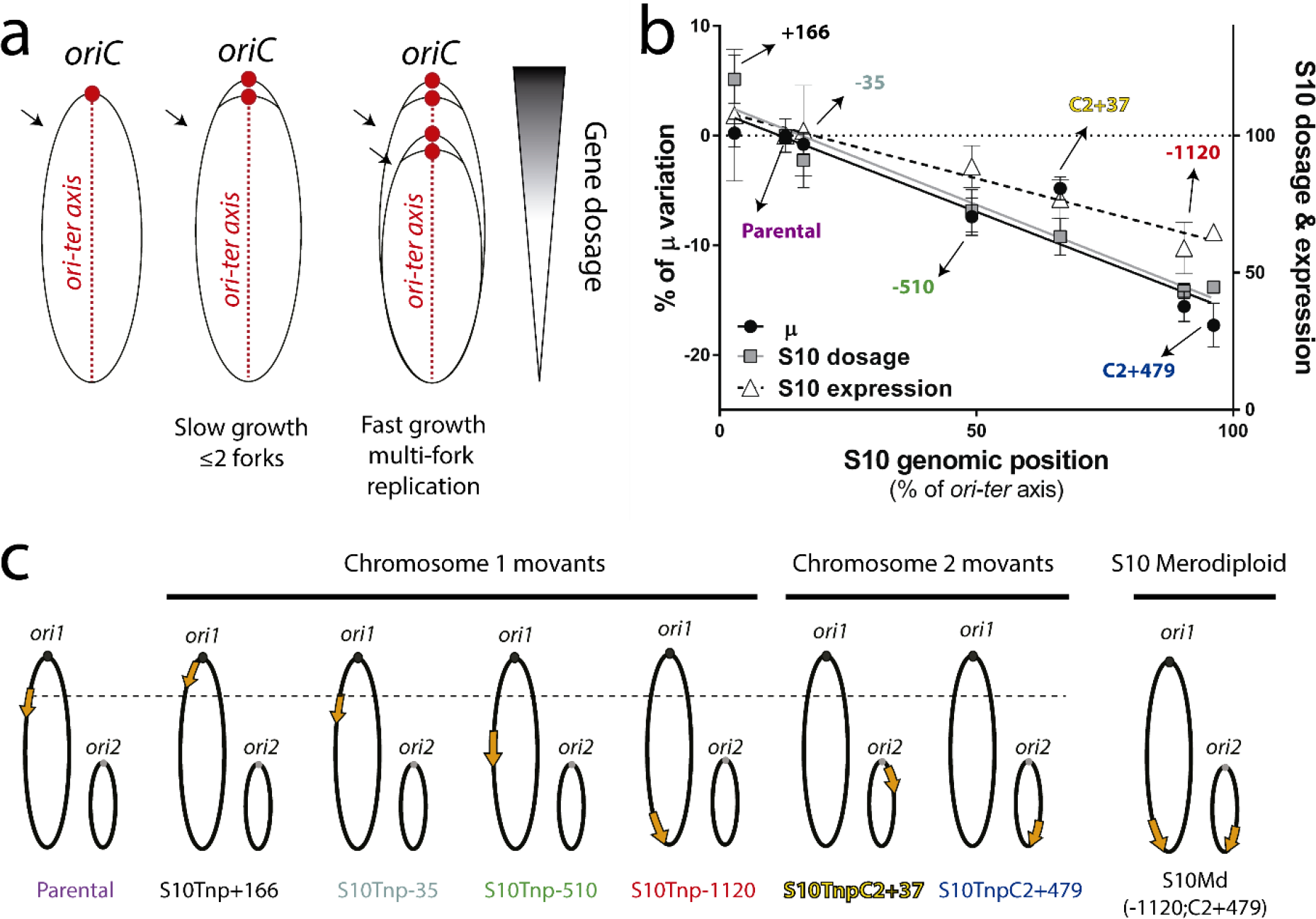
Genome organization links S10 location to cell physiology. **a)** The presence of a single *oriC* (red dot) organizes the bacterial genome along an *ori-ter* axis (left panel). In slow growing conditions, genes have between 1 to 2 copies (center). During exponential phase, fast growing-bacteria overlap replication rounds increasing the dosage of *oriC*-neighboring regions (right panel). The arrow shows the approximate position of the S10 locus. **b)** The maximum growth rate (μ, black dots) and the relative S10 dosage (gray squares) and expression (white triangles) with respect to the parental strain were plotted as a function of S10 position along the *ori-ter* axis within *V. cholerae* genome. **c)** Diagram of the genome of parental, movant, and the merodiploid strains employed in this study. *ori1* and *ori2* are depicted as dark and light gray dots, respectively. The orange arrow represents S10 displaying its genomic position and ploidy. The dashed line represents the S10 location in the parental strain. Chromosomes are drawn according to their replication timing.

Notable cases are genes encoding the flux of the genetic information. In fast-growing bacteria, the genes coding for transcription and translation machineries locate near the *oriC* (16, 17). These microorganisms divide faster than the time required for genome duplication. Consequently, chromosomes trigger replication more than once before cytokinesis, overlapping successive DNA duplication rounds, a phenomenon called multi-fork replication (Fig. 1a). This leads to replication-associated gene dosage gradients along the *ori-ter* axis during exponential growth (Fig. 1a) (14). Therefore, it was proposed that the *oriC*-proximal location of ribosomal and transcription genes allows the recruitment of multi-fork replication for growth optimization purposes (5, 16, 17). Thus, the dosage and expression of the aforementioned genes peak during exponential growth phase (Fig. 1a, right) when the transcriptional activity and ribosome numbers increase by 10 and 15-fold respectively (18).

In previous works (19, 20), we tackled this issue in *Vibrio cholerae*, the causative agent of cholera disease. This microorganism is also a model for multi-chromosomal bacteria, a trait found in ~10% of these microorganisms (21). *V. cholerae* harbors a main chromosome (Chr1) of 2.96 Mbp and a 1.07 Mbp secondary replicon (Chr2). Their replication is coordinated along the cell cycle: the *oriC* of Chr2 (*ori2*) fires only after 2/3 of Chr1 duplication has elapsed, finishing the process synchronously (22, 23). *V. cholerae* is among the fastest-growing bacteria and therefore it displays particularly high replication-associated gene dosage effects (16). Its transcription and translation genes map close to the *oriC* of Chr1 (*ori1*) (19). Among them, *S10-spc-α* (S10) is a 13.4 Kbp locus harboring half of the ribosomal protein genes (RP) located 0.19 Mbp away from *ori1* (19). Using recombineering techniques, we built a set of S10 *movants* (i.e. isogenic strains where the genomic position of S10 locus is modified) to uncover interplays between the chromosomal position of the locus and cell physiology. We found that its maximum growth rate (μ) decreased as a function of the distance between S10 and *ori1* (Fig. 1b and 1c). Also, S10 genomic location impacted on *V. cholerae* fitness and infectivity (19, 20). In line with prior bioinformatics studies (16, 17), we showed that *oriC*-proximity of S10 provides optimal dosage and expression to attain the maximal growth capacity (19). We also found that S10 position impacts bacterial fitness in absence of multi-fork replication (20). This suggests that the RP gene location affects cell physiology even in slow-growing bacteria (20). In sum, our previous work and the cited examples (14) support the notion that gene order conditions cell physiology, shaping genome structure along the evolution.

However, although we proved that the current S10 genomic location maximizes *V. cholerae* fitness (19, 20), we still lack a mechanism explaining this phenomenon. Here, we addressed this issue through the most straightforward hypothesis that is S10 relocation far away from *ori1* diminishes ribosome component availability. This in turn, should reduce ribosomal activity, impacting cell physiology globally through the general impairment of protein synthesis. In this work, we quantified the global protein production in the parental strain and in the most affected derivatives (Fig. 1b and 1c). RNA and DNA deep-sequencing revealed genome-wide alterations in gene transcription and replication dynamics. Surprisingly, we found no differences in global protein production at the population level. This suggests the existence of global mechanisms linking S10 dosage to cell physiology not linked to protein biosynthesis capacity.

The intracellular milieu has a very high concentration of macromolecules that reaches 400 mg/mL in *Escherichia coli*. Consequently, the cytoplasm does not behave as an ideal solution since this large quantity of macromolecules occupies 20-30% of its volume, which is physically unavailable to other molecules. Such steric exclusion creates considerable energetic consequences, deeply impacting intracellular biochemical reactions. This phenomenon, referred to as macromolecular crowding (24, 25), has received little attention in *in vivo* systems (26, 27). Protein accounts for ~55% of the bacterial cell mass (18, 24), with RP representing one third of them (28). We hypothesized that S10 expression reduction, would lead to lower macromolecular crowding within the bacterial cytoplasm, globally affecting cell physiology (24, 26, 27). Here, we gathered evidence supporting the idea that S10 relocation mainly impacts cellular physiology of *V. cholerae* by altering cytoplasm homeocrowding (i. e. macromolecular crowding homeostasis) (24).

## Results

### S10 relocation does not cause ribosomal activity reduction at the population level

We recently settled that S10 relocation impacts cell physiology in a dosage-dependent manner (19, 20). However, how S10 dosage reduction affects cell physiology was still unknown. The most plausible explanation is that a reduction of RP levels upon S10 locus relocation affects ribosome biogenesis leading to a reduction in protein synthesis. To inquire if S10 relocation impairs protein production, we created strains expressing GFP by inserting *gfpmut3*\* (29) under a strong constitutive promoter into an innocuous intergenic space (Table S1). The direct quantification of fluorescence, allows for estimation of protein production capacity in each strain (30). First, we followed in time the optical density (OD) and the fluorescence signal of these derivatives. We estimated translation capacity by plotting fluorescence as a function of OD (Fig. 2a). Fluorescence increased exponentially as the OD incremented (R^2^>0.99, Table S2). Although the curves differed slightly between strains, there was no significant correlation between S10 genomic position and GFP production (Pearson’s Test, r=0.1, p=0.86). We next subjected cultures of these strains to flow cytometry during the early exponential phase, when S10 dosage differences among the movants are maximal. This method allows to simultaneously observe the average GFP production per cell with higher sensitivity and the distribution of fluorescence among the cells in the populations (Fig. 2b). All tested strains showed similar signal levels and the same distribution pattern. In sum, we found no link between GFP production and S10 genomic location.

**Figure 2:**
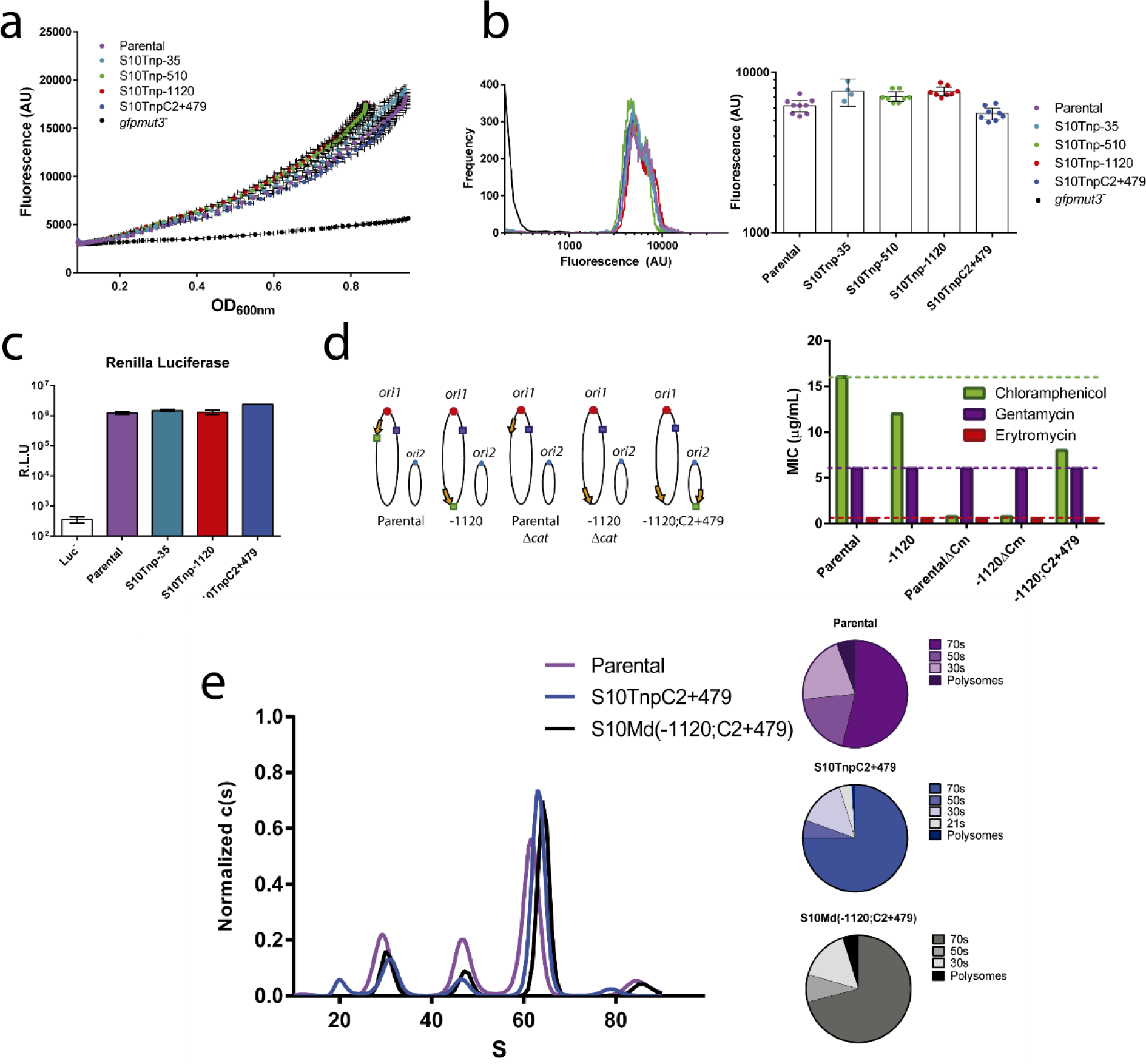
S10 genomic location does not impact ribosome function at the population level. **a)** The GFP expression and OD_600nm_ of the indicated *gfpmut3*^+^ strains (Table S1) were measured along time. The fluorescence mean (±SD) was plotted as a function of the mean (±SD) OD_600nm_. Figure shows a representative experiment with 4 biological replicates (among three independent experiments). The parental *gfpmut3*^−^ strain is an autoflourescence/light dispersion control. **b)** The indicated *gfpmut3*^+^ strains in early exponential phase (OD_450nm_~0.2) were analyzed by flow cytometry. Left panel shows the fluorescence signal frequency distribution of the indicated *V. cholerae* strains. Parental *gfpmut3*^−^ strain was added negative control. Right panel shows the Fluorescence intensity with the 95% confidence interval (CI). Points represent individual biological replicates obtained along at least 2 independent experiments **c)** Parental and movant strains bearing RLU in the chromosome (Table S1) were grown until early exponential phase. Then, RL activity, represented as RL units (RLU), was measured in three independent biological replicates for each strain. **d)** Parental and derivative strains present similar resistance levels to ribosome-targeted antibiotics. On the right panel, chromosomes are represented as in the previous figure. The encoded antibiotic resistance markers are depicted as boxes: Gm in violet and Cm in green. Their approximate genomic location is shown in each strain. On the right the MIC (μg/mL) for Cm, Gm and Er for each depicted strain is shown. e) Ribosome profiles for the indicated strains as obtained by analytical ultracentrifugation. Pie charts quantify polysome, 70s, 50s and 30 s fractions for the indicated strains.

To confirm that these results were not due to lack of sensitivity, we used the *Renilla* Luciferase (RL) as a reporter of protein synthesis capacity. RL detection shows higher sensitivity than GFP due to lower background, higher signal amplification and a larger dynamic range, making it suitable to reveal more subtle differences otherwise impossible to differentiate (31). We built S10 movant strains constitutively expressing RL at high levels (Table S1). Again, no differences in luciferase activity arose between the parental strain, S10Tnp-35, S10Tnp-1120 and S10TnpC2+479 (Fig. 2c), suggesting similar translation capacity at the population level.

As an alternative approach to look for differences on ribosomal activity, we measured the minimum inhibitory concentration (MIC) of ribosome-targeting antibiotics such as chloramphenicol (Cm), gentamicin (Gm) and erythromycin (Er). A reduction in the number of ribosomes increases sensitivity to these antibiotics (32). We measured MIC for Cm, Gm and Er using E-tests (Fig. 1d). All generated mutants derive from a *V. cholerae* strain sensitive to Er and harboring Gm resistance gene (Table S1). Strains that only differed in the genomic location of S10, had their growth inhibited at the same Er and Gm concentrations (Fig. 2d) suggesting no differences in ribosomal numbers. In parallel, the parental, S10Tnp-1120 and the S10Md(−1120;C2+479) strains harbor the Cm resistance gene (*cat*) linked to the S10 locus, therefore the location of the resistance gene differed among them (Fig. 2d). Cm resistance was higher in the Parental strain when *cat* is closer to the *ori1* and lower in S10Tnp-1120 and S10Md(−1120;C2+479) when the resistance marker is nearby the *ter1* region. Hence, as in other genetic systems (33), Cm sensitivity varied according to *cat* genomic location independently of S10 copy number (compare S10Tnp-1120 to S10Md(−1120;C2+479)). Therefore, even though this assay is sensitive enough to capture the effects caused by differences in *cat* location, it showed no antibiotic susceptibility differences related to S10 dosage. The lack of effects of S10 relocation on MIC when using any of the three different ribosome-targeting antibiotics, possessing different tolerance levels, suggests that the number of ribosomes is not affected by the genomic location of S10.

### S10 genomic location causes changes in GFP synthesis capacity at the single cell level

Since we did not detect differences in ribosomal activity at the population level, we measured GFP production at the single cell level using Fluorescence Recovery After Photobleaching (FRAP). In this assay individual cells expressing *gfpmut3*\* were photo-bleached and followed over time for at least 5 minutes. Then, we quantified the percentage of fluorescence recovery. In the parental strain, ~95% of the cells displayed a recovery of at least 20% (mean=53.8%, n=108) of the initial signal after 3 minutes, to reach a plateau until the end of the observation (Fig. S1a). The addition of Cm up to the MIC inhibited the fluorescence increase (mean=15.8%, n=21), suggesting that signal recovery corresponds to GFP re-synthesis. Meanwhile, we observed lower average recovery in the most physiologically affected movants S10Tnp-1120 (20.1%, n=42) and S10TnpC2+479 (25.8%, n=82), Fig. S1b) suggesting that they produced less GFP. Therefore, at the single cell level, the parental strain displayed a higher protein synthesis capacity than the most affected S10 movants.

### S10 relocation alters the ribosomal sedimentation profile

Reduction in RP expression can lead to problems in ribosome assembly due to modifications in the stoichiometry of its components. To detect alterations in ribosome assembly, reflected in changes in ribosomal subunits composition, we performed ribosome preparations followed by analytical ultracentrifugation (AUC) in the parental and the physiologically impaired S10TnpC2+479 strain. We also analyzed a merodiploid strain where most of the growth deficiency is rescued but still display a reduced μ (S10Md(−1120;C2+479)) (19). We expected that growth impairment would correlate with a reduction in the proportion of assembled ribosomes (i. e. the 70s peak), when compared to free ribosomal subunits (30s and 50s peaks). Figure 2e shows that parental strain displayed a 53,97% of the signal in the peak corresponding to the 70s while 50s and 30s peaks represented 19.4 and 20.8% respectively. In the S10TnpC2+479 movant, we observed an increase in the 70s proportion to the 75.85% of the signal while the free ribosomal subunits lowered to 5.5% and 14.8% of the signal for 50 and 30s subunits respectively. In the S10Md(−1120;C2+479) strain, showing an intermediate growth phenotype, 70s, 50s and 30s represented 71%, 8.3% and 15.8% of the signal respectively. Our data shows that a reduction in S10 expression led to an increase of the proportion of assembled ribosomes and a reduction of free ribosomal subunits. Therefore, movant strains might compensate lower S10 expression engaging more free subunits into translation. This could explain the relatively low impact of S10 relocation on translation capacity.

### Dosage reduction of S10 non-ribosomal genes does not impact cell physiology

Since reduction of protein biosynthesis upon S10 relocation was mild, we reasoned that it cannot explain the drastic changes observed in fitness and growth rate (μ). Meanwhile, S10 harbors genes not related to ribosome biogenesis: *rpoA,* the gene encoding for the α-subunit of RNA polymerase and *secY*, which encodes a sub unit of the Sec translocon (34), essential for protein export. We wondered whether dosage reduction of *rpoA* and/or *secY* could contribute to the phenotype caused by S10 relocation by provoking a reduction of the transcription rate and/or by hampering the normal protein export process. To test this, we cloned *rpoA* and *secY* on a low copy-number plasmid with inducible expression. The parental strain (Table S1, Parental) and the two most affected movants, S10Tnp-1120 and S10TnpC2+479 were transformed with either of these plasmids or the empty vector. Next, the μ of the transformed strains was determined through automated growth curves. If lower RNAP and/or translocon activity were involved in the observed phenotypes, growth rate differences between the parental and movant strains should lessen or disappear upon *rpoA* and *secY* overexpression. Results on Figure S2 show that the growth rate was significantly lower in the movants compared to the parental strain independently of the genes expressed on the plasmid vector. Since the plasmids expressing *rpoA* or *secY* did not rescue the growth defect, the impact of S10 relocation on cell physiology results from dosage reduction of RP genes within the locus.

### Transcriptome analysis of the movant strain set

Since the physiological effects of S10 relocation are due to dosage reduction of RP genes and the effects on translation were only observed at single cell level, we reasoned that alternative mechanisms must explain the effects observed at the population level. To detect genes whose transcription was affected by S10 relocation and search for metabolic pathways responding to RP dosage alterations we characterized the full transcriptome of: S10Tnp-35, the movant in which S10 was slightly moved presenting no phenotype; and the physiologically impaired strains S10Tnp-510, S10Tnp-1120 and S10TnpC2+479 (Fig. 1b). We collected the samples in fast growing conditions during exponential phase ensuring maximal S10 dosage differences, and then we compared each movant’s transcriptome to the one of the parental strain.

We first looked at the read coverage along the chromosomes, a parameter accounting for the genome-wide transcriptional activity. In fast growing conditions, we observed that the transcription of the *ori1-*region decreased as a function of the distance between S10 and *ori1* (Fig. 3a). To quantify this effect, we calculated the read coverage of the 400 Kbp flanking *ori1* (35). While S10Tnp-35 displays no significant transcriptional alteration within this genomic region, a significant reduction was observed in S10Tnp-510 (−1.042 fold change, p<10^−13^), S10Tnp-1120 (−1.056, p<10^−25^) and S10TnpC2+479 (−1.044, p<10^−8^) (Figs. 3a and S3). This was not the case for Chr2 where the *ori2* region displayed no transcriptional activity differences across the strains. The sole exception was a small increase in the transcriptional activity of the superintegron (36) in S10Tnp-1120 movant (Fig. S4). Therefore, a global, yet relatively small, reduction of transcriptional activity of the *ori1* region is observed upon relocation of S10 far from *ori1*.

**Figure 3:**
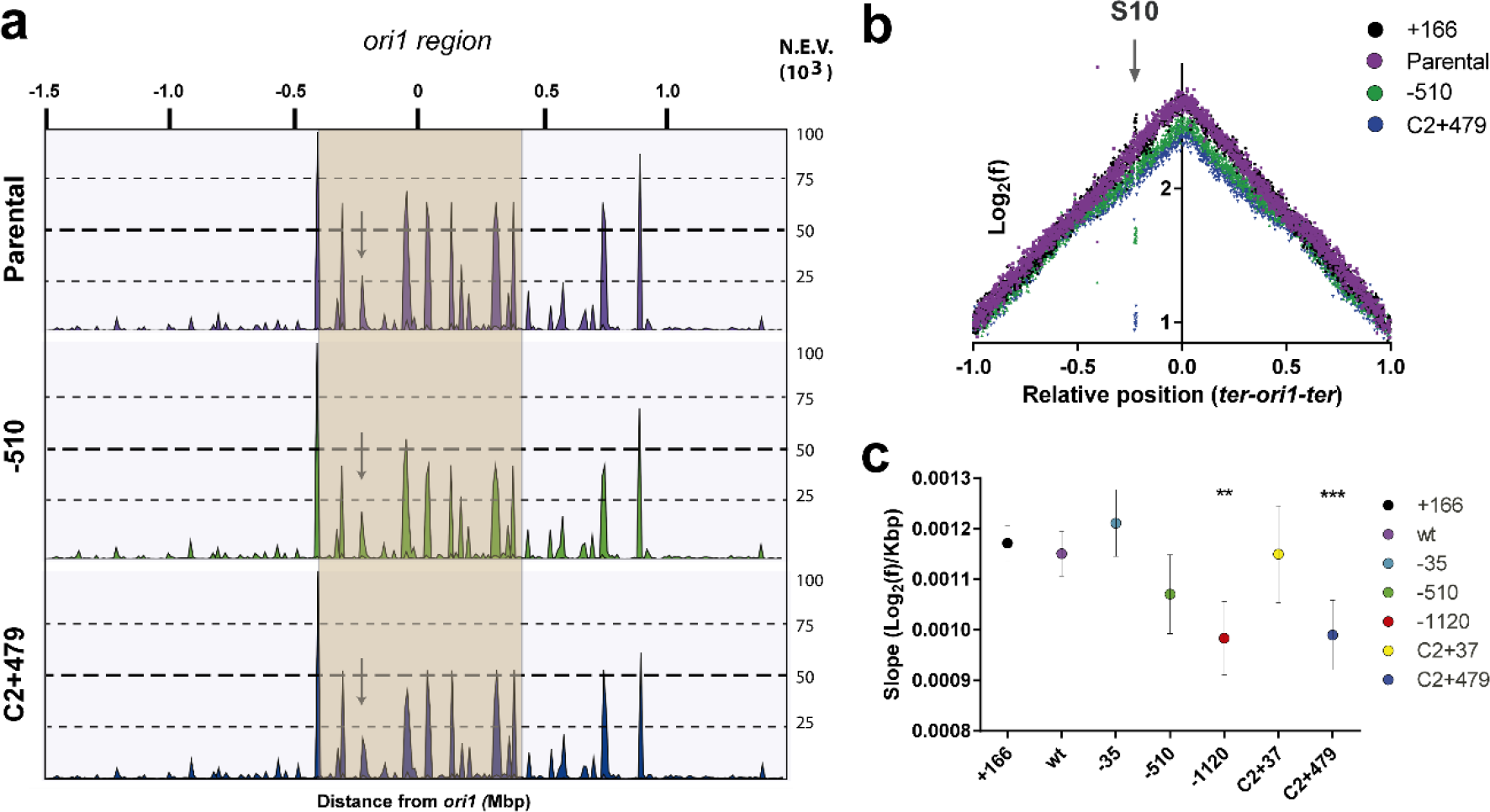
Genome-wide transcription and replication activity along the genome. **a) Transcriptional activity across Chr1.** RNA-seq reads were mapped along the Chr1 of *V. cholerae*. The histograms represent mapped read normalized to the genome wide total volume along both replichores in *ter1-ori1-ter1* order. Normalized Expression Values (NEV) are shown along the distance from *ori1* in Mbp is shown on top. Each graph represents one strain: Parental (purple); S10Tnp-510 (green); S10TnpC2+479 (blue). The plots of the whole strain set are in FigS4. The 400 Kbp flanking *ori1* are highlighted in orange. The arrow indicates the peak corresponding to the S10 locus. **b)** MFA profiles are obtained by plotting the log_2_ frequency of reads (normalized against reads from a stationary phase of a parental strain control) at each position in the genome as a function of the relative position on the *V. cholerae* main chromosome with respect to *ori1* (to reflect the bidirectional DNA replication) using 1,000-bp windows. Results for the parental (purple), S10Tnp+166 (black), the S10Tnp-510 (green) and the S10TnpC2+479 (blue) movants show their differences in read coverage. The arrow highlights the S10 position in the abscissa, reflecting dosage alterations. **c)** S10 relocation effect on replication dynamics was quantified by averaging obtained the slope for each replichore for at least 4 independent MFA experiments. Results are expressed the mean slope with 95% CI. Statistical significance was analyzed by one-way ANOVA two-tailed test. Then Tukey test was done to compare the mean values obtained for each strain. Statistically different slopes are indicated as follows: **, p<0.01 and ***, p<0.001.

### Replication dynamics are altered in the most affected movants

Given that a specific mechanism regulating the expression of such a wide genomic region seems unlikely, we wondered if the change in the expression of *ori1* region was linked to changes in global replication pattern. To assess this, we studied the replication dynamics of the genome of the whole strain set using Marker Frequency Analysis (MFA). For this, we aligned genomic DNA reads from exponentially growing cells of each strain to the *V. cholerae* genome. For each replicon, there is a linear relationship between the Log_2_ number of reads covering the locus and its genomic position between the *oriC* and the *ter* (37) (Figure 3b). This allows for robust quantification of replication dynamics across the bacterial genome with unprecedented resolution of replication fork speed and the *ori* and *ter* region locations (23, 37–39). To better quantify these differences, we calculated the average slope (Log_2_(frequency)/Kbp) along both replichores, which estimates the replication speed for each strain (Fig. 3c). MFA analysis revealed significant differences in replication dynamics across the strain set. The parental strain, the S10Tnp+166 and the S10Tnp-35 displayed a similar slope (Table S3). Conversely, the most affected movants, S10Tnp-1120 and S10TnpC2+479, where S10 was relocated at the termini of Chr1 and Chr2, showed a significantly lower slope (p<0.01, Fig. 3b, 3c and Table S3). S10Tnp-510 and S10TnpC2+37 displayed an intermediate value not significantly different from either group. Coincidentally, the calculated slope closely correlated to the S10 locus genomic position (r =-0.78, p<0.05), its dosage (r =0.8, p<0.05), the ori1/ter1 ratio (r =0.91, p<0.005) and μ (r=0.9, p<0.01) (Fig. S5). This suggests that the genomic location of S10 impacts DNA replication activity, slowing down replication when S10 is far from *ori1*. These data (Fig. 3b, 3c and Table S3) indicates that DNA coverage decreases at the *ori1* region with increasing *ori1*-S10 distance. This trend matched the changes in transcriptional coverage observed in RNA-seq data.

### Differentially expressed genes upon S10 relocation

We next analyzed the transcriptomic data to find which genes and pathways were differentially transcribed with respect to the parental strain in S10Tnp-35 and in the affected movants S10Tnp-510, S10Tnp-1120 and S10TnpC2+479 (Fig. 1b and 1c).

First, using volcano plots, we analyzed the statistical significance of the changes in transcription of each gene (−Log_10_(*p-value*)) as a function of its transcriptional Log_2_ of fold change (Log_2_(FC)) compared to the parental strain. We observed more transcriptionally altered genes with higher distances between the S10 locus and *ori1* (Fig. 4a). S10Tnp-35, a strain presenting no phenotype used as a control of the neutrality of the relocation process, displayed only 8 genes with significant (p<0.05) transcriptional change (Table 1, Data Set 1). S10Tnp-510, displaying a slight μ reduction (Fig. 1c), showed 111 genes with significantly altered transcription (Table 1, Fig. 4a, Data Set 1). Finally, the most affected movants, S10Tnp-1120 and S10TnpC2+479, displayed a significant transcriptional change in 664 and 742 genes, representing 17.95% and 20.06% of their gene repertoire, respectively. Most of altered genes in the movants were up regulated (Fig. 4b and Table1). These transcriptional perturbations were relatively small in magnitude since only a 26%, a 10.8% and a 14.15% of altered genes presented alterations greater than 2-fold in S10Tnp-510, S10Tnp-1120 and S10TnpC2+479 respectively. Meanwhile, up-regulated genes showed 2.8-fold, 1.6-fold and 1.7-fold average increases respectively (Table 1, Fig. 4b, Data Set 1). In the three movants, the down-regulated genes displayed a smaller perturbation of ~1.4-fold (Table 1).

**Table 1:** Quantitative and qualitative expression changes in the movant strains. The number of differentially expressed genes (p<0.05) compared to parental strain in fast growing conditions. The number in parenthesis represents genes whose expression varies more than 2-fold. The magnitude of expression change is quantified as the average of the Log_2_(FC) ± standard deviation.

|  | **-35** | **-510** | **-1120** | **C2+479** |
| --- | --- | --- | --- | --- |
| **Number of upregulated genes** | 2 (1) | 62 (37) | 361 (64) | 439 (88) |
| **Mean upregulation<sup>a</sup>** | n/d | $1.5 \pm 0.97$ | $0.67 \pm 0.41$ | $0.78 \pm 0.56$ |
| **Number of downregulated genes** | 6 (4) | 49 (2) | 301 (9) | 303 (17) |
| **Mean downregulation<sup>a</sup>** | n/d | $-0.5 \pm 0.24$ | $-0.49 \pm 0.26$ | $-0.52 \pm 0.29$ |
| **Total number of altered genes** | 8 | 111 (39) | 662 (72) | 742 (105) |
| **Altered functions** | - | E, P, V | E, O, R, V, N | E, O, R, V, N, F, P, U |

<sup>a</sup> average of the $\text{Log}_2(\text{FC}) \pm$ standard deviation. n/d, not determined.

**Figure 4:**
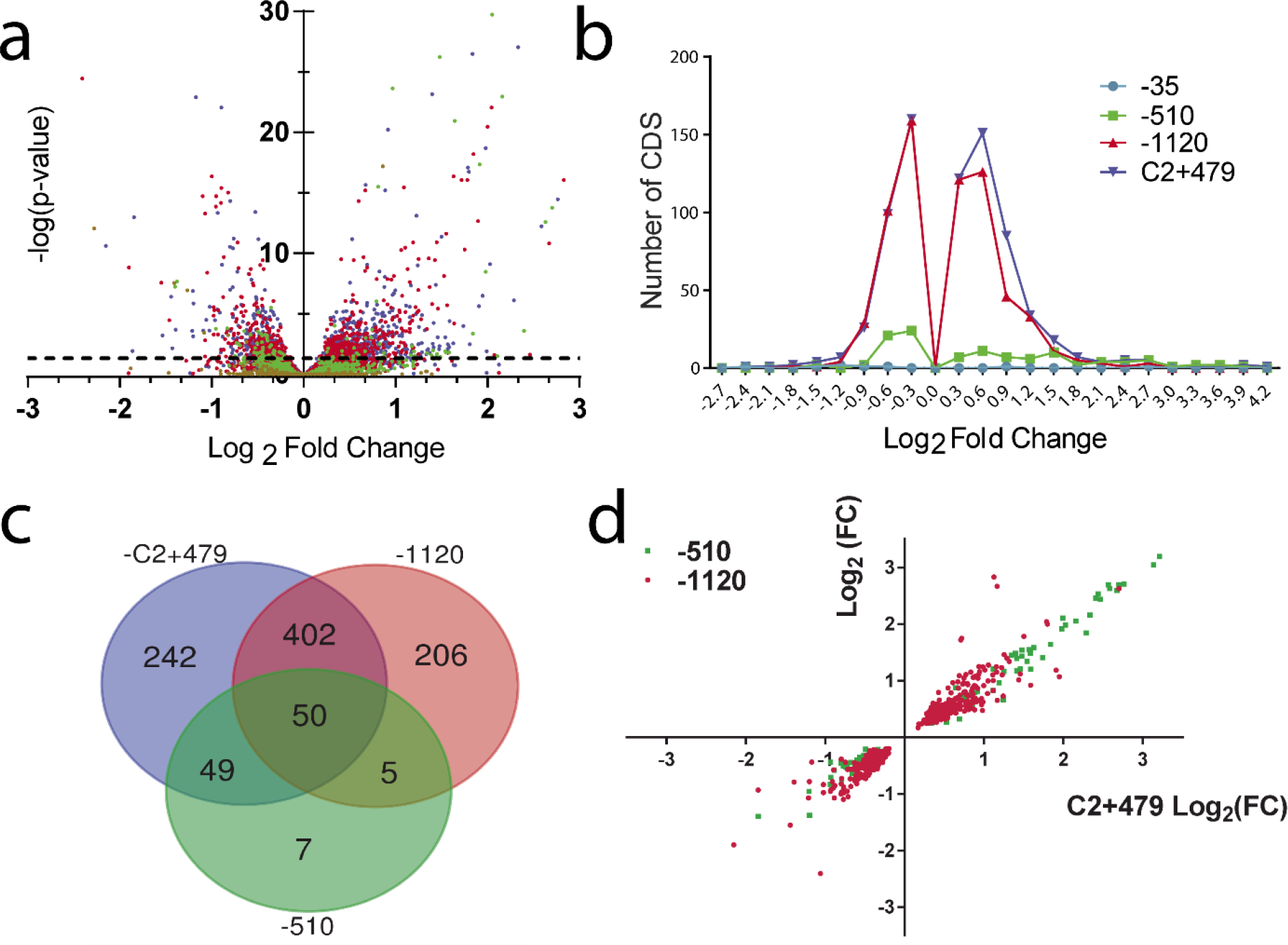
S10 relocation impacts gene expression genome-wide in a distance dependent manner. **a)** Volcano plot displaying differential expressed genes in S10Tnp-35 (brown), S10Tnp-510 (green), S10Tnp-1120 (red) and S10TnpC2+479 (blue). Horizontal dotted line shows p=0.05. **b)** The number of coding sequences (CDS) as a function of Log_2_(FC) of strains S10Tnp-35 (turquoise), S10Tnp-510 (green), S10Tnp-1120 (red) and S10TnpC2+479 (blue). **c)** Venn diagram displaying shared genes between S10Tnp-510 (green), S10Tnp-1120 (red) and S10TnpC2+479 (blue). **d)** Expression correlation between movant strains. Dots correspond to individual CDS. The Log_2_(FC) of each gene in S10Tnp-510 (green) or S10Tnp-1120 (red) was plotted as a function of Log_2_(FC) in S10TnpC2+479.

Most of the transcriptional alterations were found in the same genes across the S10 movants (Figure 4c). A large fraction of transcriptionally altered genes in a movant were also regulated in either of the other two movants (Table S4). Shared genes showed similar levels of transcriptional change across the movants (Fig 4d and Table S4). For example, the degree of change in altered genes of S10Tnp-510 and S10Tnp-1120 were highly correlated (r=0.927, p<10^−24^). The differentially expressed genes were not confined to specific chromosome regions nor associated to a specific replicon: S10 relocation produced homogeneously distributed changes in *V. cholerae* gene transcription (Fig. S6).

To identify the functions or metabolic pathways altered by S10 relocation, we classified *V. cholerae* genes in 25 functional categories using the <u>EMBL</u> eggNOG database v.4.0 (40)(Supp. Text). We then identified the categories with over or under-representation of genes with altered transcription levels in S10Tnp-510, S10Tnp-1120 and S10TnpC2+479 with respect the full repertoire of *V. cholerae* genome (Data Set 1, Table S5, Fig. S7)

Genes from the category ‘Translation, ribosomal structure and biogenesis’ (J) were not significantly altered, which is consistent with the results above showing that S10 relocation did not alter the translation capacity (Fig. 2). The category ‘Amino acid transport and metabolism’ (E) was statistically altered in all three movants. The category “Posttranslational modification, protein turnover, chaperones” (O) was the most affected category in S10Tnp-1120 and S10TnpC2+479, since about 65% of its genes showed higher transcription in the movants (Table S5, Data Set 1). The list of up-regulated genes was dominated by chaperones and heat-shock proteins. Strikingly, the highest transcriptional changes occurred in the main pathway for cytosolic protein folding (41): *grpE* (VC0854), *dnaKJ*(VC0855-6) and both copies of the *groEL-groES* system (VC2664-5 and VCA0819-20). Many transcriptionally altered genes were involved in protein export and ion transport, belonging to several significantly perturbed categories such as: “V, Defense mechanisms” (e.g. VC0590 coding for an ABC-2 type transporter), “U, Intracellular trafficking, secretion, and vesicular transport” (*secA,* VC2462), and “N, cell motility” (some *fli* and *fla* genes) (Table 1 and Data Set 1). Some particularly induced genes of “P, Inorganic ion transport and metabolism” group were iron (*hutX, hmuV, hmuU, exbD1, tonB;* ~2.8 FC) and sulfur (*sbp*, *cysHI, cysDNC;* ~8 FC) transporters. Based on the analysis of functional categories, we observed that *V. cholerae* responds to S10 relocation by altering amino acid synthesis pathways, increasing the transcription of chaperones and proteases probably to degrade misfolded proteins and by activating the expression of transporters and permeases.

### Cytoplasm is more fluid in the most affected movants

During exponential growth, ribosomes account for up to 30% of bacterial dry weight (42). S10 encodes half of the ribosomal proteins, which are very highly expressed constituting more than a third of total cell proteins in *E. coli* (28). Therefore, it is likely that a reduction in S10 expression results in macromolecular crowding alterations as observed in other systems (43, 44). Macromolecular crowding is crucially important in biochemical reactions, however how it impacts cellular physiology remains mostly unexplored (24–26). It is well documented that it influences protein folding, aggregation and perturbs protein-nucleic acids interactions (45). On the other hand, DNA replication has an absolute dependence on macromolecular crowding (44, 46). Therefore, the reduction in replication fork dynamics (Figs. 3b and c), the alteration of genes linked to protein folding, protein degradation, permeases and transport systems (Data Set 1, Table 1) observed upon S10 relocation can be interpreted in light of changes in macromolecular crowding caused by a lower RP concentration.

To test this hypothesis, we measured the viscosity of the cytoplasm in the parental strain and in the most affected movants, S10Tnp-1120 and S10TnpC2+479. We expected a more viscous cytoplasm in the parental strain since it expresses S10 genes at higher levels generating a greater concentration of RPs than the movant strains. Differences in cytoplasm viscosity can be uncovered by FRAP experiments on GFP expressing strains. For this, the fluorescence recovery time is measured after bleaching a part of the bacterial cytoplasm (47, 48). Since the small size and the comma-shape of *V. cholerae* complicates the procedure, we generated elongated cells by deleting the Chr2 replication-triggering site (*crt*S) (23) in cells expressing GFP. These mutants present a defective replication of the secondary chromosome. Therefore, S10TnpC2+479 should have even less copies of S10 per cell and, concomitantly, display higher cytoplasmic fluidity than S10Tnp-1120.

In the *gfpmut3\** Δ*crtS* context (Table S1), the parental strain displayed a significantly longer half-time recovery of fluorescence (τ) than the movants (Fig. 5a, Supp. Text). The collected data showed a high dispersion due to biological variability, however, τ distribution was different in the movants when compared to the parental strain (Fig. 5b) which displayed a τ of 139.7 ms (95% confidence interval (CI) (120.4-158.9)ms; median=110 ms; n=104). As expected, S10Tnp-1120 showed a τ of 97.3 ms (95% CI (88.31-106.3)ms; median=90 ms; n=128) significantly shorter than the parental strain (p<0.0001). S10TnpC2+479 displayed a τ of 107.5 ms (95% CI (97.39-117.52) ms; median= 100 ms; n= 92), statistically lower than the parental strain (p<0.05) but not significantly different from S10Tnp-1120. The more fluid cytoplasm in movants could be a consequence of fewer S10-encoded RP suggesting that S10 relocation far from *ori1* reduces cytoplasm macromolecular crowding.

**Figure 5:**
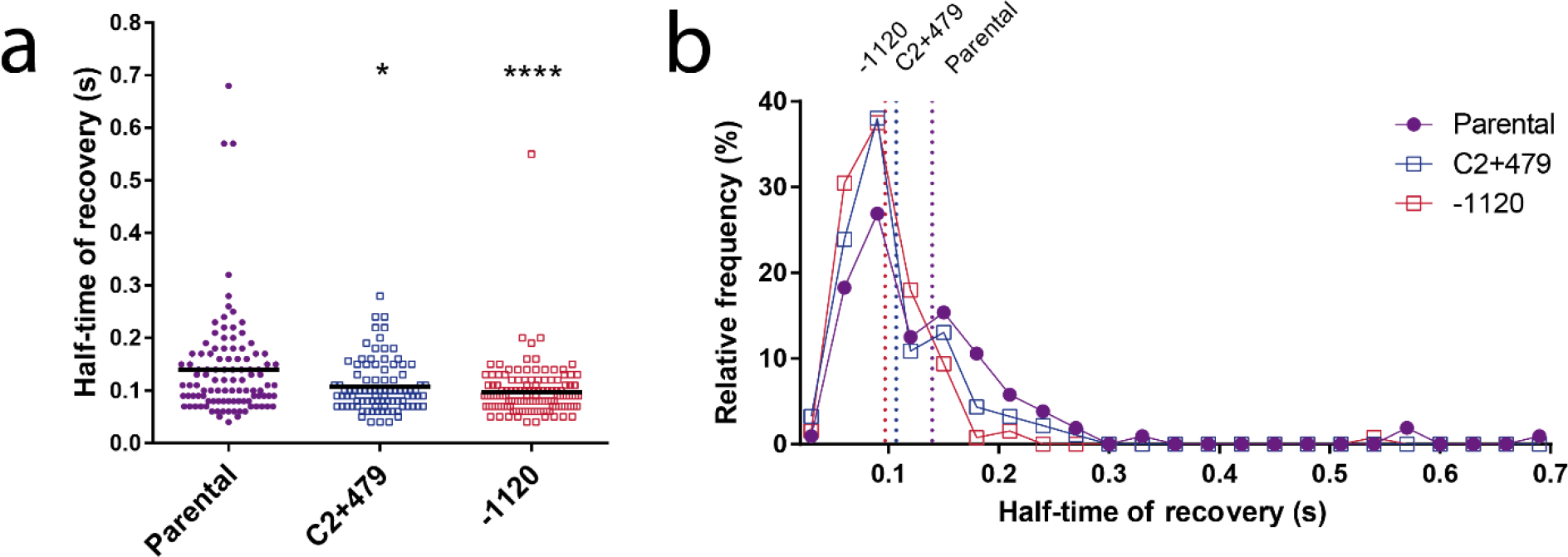
S10 relocation impacts cytoplasm fluidity. **a)** Half-time of fluorescence recovery (τ) in the Parental-1120 (purple, n=104) and the most affected movants S10Tnp-1120 (red, n=128) and S10TnpC2+479 (blue, n=92) in a *gfpmut3\** Δ*crtS* genetic context. The line indicates the mean τ value and each dot indicates the obtained value for a cell. Statistical significance was analyzed using Kruskal-Wallis non-parametric tests followed by Dunn’s multiple comparisons using parental as control respectively. *, p<0.05; ****, p<0.0001. **b)** Histogram showing the relative frequency of τ to observe the distribution of the values. The vertical dotted line shows the mean value as in **a)**.

### Growth rate and replication dynamics alterations in movants are alleviated in hyperosmotic conditions

In line with lower macromolecular crowding, we observed a reduction in cytoplasm viscosity in the movants. To test the possible impact of such molecular crowding alterations on the physiology of the movants, we employed an osmotic stress approach (49–51). This consists of subjecting strains to a hyperosmotic environment. In these culture conditions, water exits the cell reducing the macromolecular crowding differences between the strains. Therefore, μ differences between the parental strain and the movants should be reduced with increasing solute concentration. To test this, we performed automated growth curves in rich media with increasing NaCl concentrations, comparing the μ of the parental strain to S10Tnp-1120 and S10TnpC2+479 movants. As depicted in Figure 6a, growth rate differences between the parental strain and the movants were reduced as NaCl concentration increased. Since this phenomenon could be explained by the nature of the solute of choice (e.g. putative differential sensitivity to NaCl), we repeated these assays using sucrose as an alternative compound. As shown in Figure 6b, results were very similar, suggesting that this phenomenon depends on osmotic changes and cannot be attributed to the nature of the solute. Notably, the μ of the parental strain was not significantly reduced in the range of 5 to 20 gr/L NaCl (Fig. S8). Meanwhile, the growth of movant strains varied significantly along this concentration range, displaying a reduced growth at 5 gr/L and 10 gr/L and reaching its maximum at 20gr/L (Fig. S8). Consequently, growth differences observed are not due to impairment of the parental strain in hyperosmotic conditions. We conclude that μ differences caused by S10 relocation far from *ori1* can be counterbalanced by artificially increasing cytoplasmic crowding.

**Figure 6:**
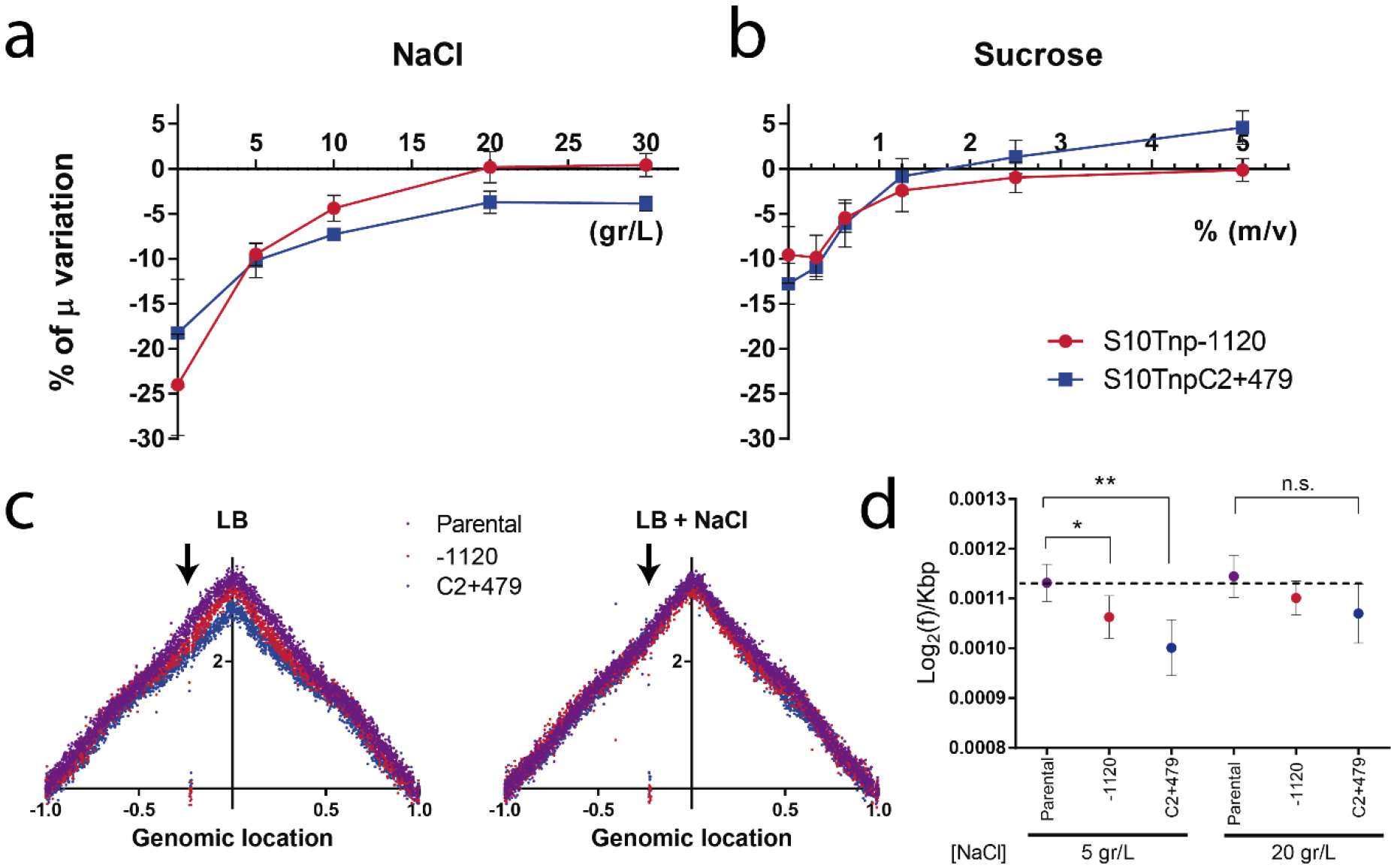
S10 relocation effects are reduced in hyperosmotic conditions. **a)** Growth rates of the parental and the indicated movant strains in LB with increasing NaCl concentrations were quantified by averaging obtained μ in 6 independent experiments with at least 3 biological replicates. The growth of each movant was normalized to the μ of the parental strain and the percentage of the variation (μ %) ± SEM with respect to parental strains is shown as a function of NaCl concentration of growth medium. **b)** Changes in growth of the movant strains with respect to parental strain is shown as a function of sucrose concentration. Data was trated as in a) but results correspond to 4 independent experiments with at least 3 biological replicates. **c)** MFA profiles are plotted as in Fig. 3b. Results for the parental (purple), the S10Tnp-1120 (red) and the S10TnpC2+479 (blue) strains in LB in presence of 5 gr/L (LB, left panel) or 20 rg/L (LB+NaCl, right panel) are shown. The arrow highlights the S10 position in the abscissa, reflecting S10 dosage alterations. **d)** Replication dynamics in presence of 5 or 20 gr/L of NaCl assessed by calculating the slope for each replichore for 2 independent MFA experiments. Dots indicate mean ± SD. Statistical significance was analyzed by one-way ANOVA two-tailed test and Tukey test for multiple comparisons. Significance is indicated as follows: n.s.: non-significant; *, p<0.05 and **, p<0.01.

Upon S10 relocation far from *ori1*, we observed a lower replication speed in the movants suggesting that DNA replication activity diminished, suggesting a lower replication speed in the movants (Fig. 3c). Since, molecular crowding is crucial for chromosome replication (44, 46) we used the osmotic stress approach to test if the observed replication dynamics defects in movants could be compensated. For this, we performed MFA analyses of the parental strain and the S10Tnp-1120 and S10TnpC2+479 movants in the presence of 5 or 20 gr/L of NaCl. In these culture conditions, the parental μ is unaffected. In contrast, movant strains grew 10-15% slower than the parental strain but they were able to rescue the growth defect at higher NaCl concentrations (Figs 6a and S8). As in earlier experiments, MFA analyses revealed that the movants have a significantly lower slope than the parental strain. Increasing NaCl concentration to 20 gr/L made their slopes converge diminishing replication dynamics differences (Fig. 6c and 6d). The integration of these and the previous observations, suggests that lower expression of RP caused by S10 relocation (Fig. 1b) leads to lower molecular crowding (Fig. 5), which negatively impacts replication (Fig. 3b). This fits the observation that addition of external NaCl, causing water loss and thus narrowing differences in macromolecular crowding, produces more similar replication dynamics between the parental and the movant strains (Fig. 6d).

## Discussion

Comparative genomics suggests that gene order coordinates cell cycle to the expression of key functions necessary for cellular homeostasis (4, 11, 16, 17) but few papers provided experimental support (13, 14, 52). A notable case is that of ribosomal genes which are located near the *oriC* in fast growing bacteria (16, 17). By systematically relocating S10, the main cluster of RP genes (Fig. 1c), we proved that its genomic location determines its dosage and expression in *V. cholerae* (Fig. 1b). S10 repositioning far from *ori1* leads to larger generation times, lower fitness and less infectivity (19, 20). These effects are dependent on S10 dosage. However, the mechanism explaining how RP dosage affects cell physiology was still missing. The most straightforward explanation was that high RP dosage due to multi-fork replication increases their expression maximizing protein biosynthesis capacity (16, 17). Our initial hypothesis was that movants in which S10 was far from *ori1* would have a lower translation capacity, easily explaining lower growth and fitness of these movants. Surprisingly, we found that in the most affected movants, translation capacity reduction could not explain the observed physiological changes (Fig. 2). We do not rule out that translation impairment may have an effect in the cellular physiology, however, it must have a secondary role in the phenotypes displayed in the affected movants. Slight differences in protein production between the parental strain and the most affected movants could only be detected at single cell level (Fig. S1). The movants displayed a larger proportion of assembled ribosomal subunits. This might compensate putative deficiencies in the translation apparatus (Fig. 2e). Interestingly, the S10TnpC2+479 displayed a small peak of ~21s that might correspond to precursors of 30s subunit typically associated to cells displaying ribosome assembly deficiencies (53). Meanwhile, complementation of movants with *secY* and *rpoA*, two S10 genes not related to ribosome biogenesis, failed to rescue the growth defect demonstrating the relevance of RP in the observed phenotype. In sum, although dosage reduction of S10-encoded RP genes caused the observed phenotypes, it is unlikely that this is a consequence of translation defects.

Deep sequencing techniques revealed less transcriptional activity in the region flanking *ori1* (Fig. 3a) and lower replication velocity in the most affected movants (Figs. 3b, 3c and 6c). Since highly expressed genes that account for a large majority of transcriptional activity in the cell (i.e. *rrn*, ribosomal protein genes, etc.) cluster at this chromosomal region, slight changes in its dosage may globally impact cell physiology (4, 11) and may be responsible for the slight reduction in translational activity observed at single cell level (Fig. S1). Meanwhile, differential expression analysis revealed that the transcriptional response is not limited to the *ori1* region (Fig. S6), and encompasses a large number of genes that show slightly but consistently altered transcription in the most affected movants (Fig 4). Furthermore, the number of these genes increases with distance between S10 and *ori1* (Table 1, Fig. 4a, 4b and S6). The latter observation corresponds to biologically meaningful transcriptional changes since furthest relocations caused larger perturbations (Figs. 4a and 4b), the majority of altered genes were common to the different movants (Fig 4c), where they showed similar transcriptional changes (Fig. 4d). This strongly suggests the presence of a common mechanism that slightly affects gene expression at a large scale. Amino acid metabolism and transport genes were less transcribed while there was an up-regulation of genes helping protein folding and cellular transporters (Table S5, Data set 1). Importantly, and in line with previous data (Fig. 2), the transcription of translation genes seems to be unaffected in the movants reinforcing the notion that lower protein biosynthesis capacity was not enough to explain the physiological alterations that we observed. Molecular crowding has a well-known key role in biochemical reactions. Even if its impact on physiological processes has been poorly studied (26), two processes - DNA replication and protein folding - are strongly influenced by macromolecular crowding (27). Since the discovery of DNA replication, the presence of crowding agents such as polyethylene glycol was shown to be absolutely necessary to reproduce DNA polymerase activity *in vitro* (44, 46). In parallel, macromolecular crowding greatly impacts protein aggregation and folding (27), although the *in vivo* consequences of how the latter occurs are still a matter of debate (45, 54). It was recently shown that ribosomes are important contributors of macromolecular crowding in the cytoplasm both in prokaryotic and eukaryotic systems (43, 44). All this information leads us to suggest that upon S10 relocation, the consequent fewer RP may lead to homeocrowding (24) perturbations. Interestingly, to the best of our knowledge this is first study exploring the consequences of lower macromolecular crowding conditions since most works linking this physicochemical factor to physiology focus on situations of increased crowding (44, 55, 56). Concomitantly, we observed reduced replication activity (Fig. 3c), as well as induction of proteases and chaperones to cope with protein aggregation and misfolding (Table 1 and Fig. S6). Notably, in the most affected movants, the genes coding for the three main chaperone systems –*grpE, dnaKJ* and *groEL-groES* (41)-were among the most strongly induced. The lower transcription of protein and ion transporters could be used for intracellular environment restoration (Table S4, Fig. S6) and could be a natural consequence of the change in cytoplasm osmotic pressure. We next tested experimentally if S10 relocation could alter homeocrowding. First, using FRAP, we observed slight but statistically significant alterations in the fluidity of the cytoplasm of the most affected movants compared to the parental strain (Figs. 5a and 5b, Supplementary Text). This supports the notion that lower expression of RP associated with movants lowers cytoplasm macromolecular crowding. In the *ΔcrtS* context, we did not detect differences in cytoplasmic fluidity between the S10Tnp-1120 and S10TnpC2+479 movants, expected from lower S10 copy number in the latter by Chr2 loss. We believe that the detrimental effects of *crtS* deletion (23) can explain this. In the S10TnpC2+479 movant, S10 dosage reduction enhances fitness loss, as reflected by slower growth and the presence of small non-viable cells in the microscope not further analyzed (data not shown). When Chr2 replication is inhibited, the fusion of both chromosomes-mainly between their terminal regions-occurs at relatively high frequency (57). Therefore, the S10TnpC2+479 *ΔcrtS* population might in part consist of cells with fused chromosomes. In this scenario S10 dosage would not decrease below 1 copy per cell.

The osmotic shock approach provided strong evidence supporting the notion that S10 dosage deficit perturbs cellular homeocrowding. In rich medium, movant strains grow slower than the parental strain. With increasing solute concentrations this growth deficit is reduced (Figs. 6a and 6b). In the case of NaCl, the parental strain grew normally in the range from 5 to 20 gr/L. Outside of this range, growth rate was reduced. Growth was particularly impaired at concentrations below 5 gr/L where culture development was very variable due to hyposmotic stress (Fig. 5b and Data not shown). Interestingly, movants looked more sensitive than the parental strain to lower solute concentrations. We think that movants express less ribosomal proteins which account for a large fraction of the bacterial proteome, which in turn constitutes a large proportion of the cytoplasmic macromolecules (58). It is known that about 0.5 gr of water is bound per gram of cytoplasmic macromolecules (49, 59). Therefore movants may lose their capacity to retain water, suffering from a situation similar to being exposed to hyposmotic conditions. Meanwhile, the μ of the parental and the movants was similar when exposed to 30 gr/L of NaCl. This indicates that detrimental hyperosmotic conditions altered the strains similarly.

Recent work shows that specific ribosomal protein genes link cell growth to replication in *Bacillus subtilis* (60). We observed similar effects since S10 dosage correlated growth rate and *oriC*-firing frequency (Fig. 3b, 3c, S6 and Table S3). In the cited study, the authors attribute this effect to ribosomal function. Although in our system the effects were milder, we do not rule out the possibility that S10 relocation alters cellular physiology through a reduction in protein synthesis. But this effect is unlikely to account for the full magnitude of the observed phenotypes (Fig. 2) especially as it is relieved in hyperosmotic conditions. We believe that this could be due to a number of factors including: i) the many regulatory mechanisms that control ribosomal protein expression at the translation level, which could partially compensate transcription reduction; ii) the fact that ribosomal subunits are found in excess with respect to assembled ribosomes; iii) the possibility that an eventual reduction in functional ribosomes can be compensated by faster translation rates (61–63); iv) finally, it has been described, particularly in *Vibrio* sp. CCUG 15956 (64), that ribosomes are available in excess of numbers needed for exponential growth. Such large ribosome quantities would have been selected as an ecological survival strategy that allows for fast growth restoration after its arrest in rapidly changing environmental conditions (65). Hence, lower S10 expression could be buffered at many levels and protein production might be only mildly impacted. Molecular crowding reduction might however not be as easily compensated. Therefore, movant strains possess a less crowded cytoplasm where DNA polymerase activity is reduced and more chaperones are needed. This would embody a novel mechanism which could explain how ribosomal protein gene position influences growth rate.

Bacterial growth closely correlates to ribosomal protein content. This has been attributed to the role ribosomes have in protein synthesis (66, 67). We propose that, on top of that, ribosome concentration may change the macromolecular crowding conditions to optimize biochemical reactions, in particular in protein folding and DNA replication (26, 27). We provide evidence indicating that this is the case for replication dynamics in *V. cholerae*. Our experiments suggest that the genomic position of S10 contributes to generate the RP levels necessary to attain optimal cytoplasmic macromolecular crowding. Besides connecting ribosomal gene position to growth in *V. cholerae*, this mechanism could link ribosome biogenesis to cell cycle in bacteria. During exponential phase, when RP production is maximal and ribosomes represent 30 % of cell weight, crowding peaks. This leads to the highest *oriC*-firing frequency. Upon nutrient exhaustion, ribosome production is reduced, the cytoplasm macromolecular crowding diminishes, slowing down replisome dynamics. This scenario, which is beyond the scope of our study, deserves to be tested in other model microorganisms.

## Materials and methods

### General procedures

Genomic DNA was extracted using the GeneJET Genomic DNA Purification Kit while plasmid DNA was extracted using the GeneJET Plasmid Miniprep Kit (Thermo Scientific). PCR assays were performed using Phusion High-Fidelity PCR Master Mix (Thermo Scientific). Strains and plasmids used in this study are listed in Table S1. Details of culture conditions and selection can be found in Supp. Text.

### Automated growth curve measurements

ON cultures were diluted 1/1000 in LB. Bacterial preparations were distributed at least by triplicate in p96 microplates. Growth-curve experiments were performed using a TECAN Infinite Sunrise microplate reader, following the OD_600nm_ every 5 minutes at 37°C on maximum agitation. Growth rate was obtained using a custom Python script coupled to the Growthrates program (68).

### Protein production capacity

For estimating GFP production we performed *V. cholerae gfpmut3*\* automated growth curves in a TECAN Infinite 200 microplate reader, following OD_600nm_ and GFP fluorescence over time. Data was analyzed using GraphPad Prism 6. For flow cytometry strains were grown in fast growing conditions until early exponential phase (OD_450_~0.2). Then 50 μL were diluted in 800 μL of PBS. The fluorescence of 20.000 events was recorded in a MACSQuant 10 analyzer (Miltenyi Biotec). Cells were detected using Side Scatter Chanel (SSC) in log_10_ scale. Data analysis was done using Flowing Software 2.5.1 (www.flowingsoftware.com). For luciferase activity measurement, *Vibrio cholerae::RL* strains were cultured until OD_450nm_~0.2. For each experiment, three samples of 20 μL were harvested and directly measured using the Renilla Luciferase Assay System (Promega).

### Ribosome profiling

Ribosomal 70s, 50s and 30s species from the indicated *V. cholerae* strains were isolated as previously described (69, 70). Early exponential phase cultures (OD_450nm_~0.2) were harvested by centrifugation. Subsequent steps were performed at 4°C. The pellet was resuspended in ice-cold Buffer A (20 mM HEPES pH 7.5, 50 mM NH4Cl, 10 mM MgCl2, 5 mM β-mercaptoethanol, 0.1 mM PMSF) in the presence of Ribolock (Thermo Fisher Scientific). DNase I was added up to 2 μg/mL and kept for 20 min at 4°C. Cells were lysed by two passes at 11,000-15.000 psi using Emulsiflex. Cell debris were removed by two centrifugation steps at 30,000*g* for 30 min. Then 0.8 mL of Cold 60% sucrose buffer A was added to RNAse-free 5 mL Ultraclean tubes for ultracentrifugation in a SW55Ti (Beckman). The ribosome-containing supernatant was used to fill these tubes and an ultracentrifugation step was performed for 16 hs at 150.000*g*. Ribosomes were recovered from the bottom 0.8 mL of 60% sucrose Buffer A and dialyzed using a Float-a-lyzer G2 in Buffer A. Sedimentation velocity was determined in a Beckman XL-I Analytical Ultracentrifuge. Double sector quartz cells were loaded with 400 μl of Buffer A as reference and 380 μl of sample (3 μm), and data were collected at 120,000 rpm from 5.8 to 7.3 cm using a step size of 0.003 cm without averaging. Sedimentation velocity data were analyzed using the continuous size-distribution model employing the program SEDFIT.

### FRAP

For measurement of GFP synthesis, stationary phase cultures of *V. cholerae* strains were diluted 1/300 in fresh LB. Then 6 μL were distributed on an LB agar pad within a Gene Frame (Thermo-Fisher) and covered with a cover slip. When indicated, the agar pad was supplemented with Cm at MIC. Cells were then visualized and recorded in a Spinning-Disk UltraView VOX (Perkin-Elmer) equipped with two Hamamatsu EM-CCD (ImageEM X2) cameras. Photobleaching was done using 5-20 % of laser power. Image analysis is detailed in Supp Text.

### Transcriptomic analysis

Preparation of RNA and libraries is detailed in Supp. Text. four independent biological replicates for each sample were done for statistical analysis which is also detailed in the Supp. Text. Trimmed reads were aligned to the *V. cholerae* reference genome using Bowtie (71) with default parameters. Aligned reads were counted using HTSeq Count (72). Further quality control and differential expression analysis was performed using methods described in supplementary methods (73–75). Graphics were done using Graph Pad software, specific online service for Venn diagram (http://bioinformatics.psb.ugent.be/webtools/Venn/) and Circos Plot (76). The sequence data was submitted to the GenBank Sequence Read Archive. Accession numbers for these samples are: SRR8316520, SRR8316521, SRR8316528, SRR8316529, SRR8316526, SRR8316527, SRR8316524, SRR8316525, SRR8316522, SRR8316523, SRR8316530, SRR8316531, SRR8316518, SRR8316519, SRR8316516, SRR8316517, SRR8316514, SRR8316515, SRR8316512 and SRR8316513.

### Whole chromosome transcriptional activity comparisons

Reads were mapped as previously described (35) to a custom assembled linear version of the *V. cholerae* that starts (base 0) at the *ter* and finishes at the *ter*, with the *ori1* at the center of the sequence. Total reads mapped to this sequence were counted and normalized as previously described (35). Fold changes were calculated using normalized values and p-values were calculated as previously described (35).

### Functional characterization of the transcriptomic response

*V. cholerae* N16961 genes were aligned against the eggNOG database v.4.0 (40). Only hits with at least 50% similarity and e-value < 0.05 were used. Each protein was assigned to the best functional category, according to the percentage of similarity and the length of the alignment. We then calculated the fraction of categories enriched in the fraction of differentially expressed genes, compared to abundances of the different eggNOG categories in the *V. cholerae* genome. The over‐or under-representation of protein families was assessed statistically using the Pearson Chi square test with Benjamini–Hochberg correction for multiple test. For further validation, this test was performed 10,000 times in random sub samples of 30% of the differentially expressed genes.

### MIC determination

The MICs of Gm, Cm and Er were determined using E-test® and the disk diffusion method following manufacturer’s instructions (Biomérieux).

## Supporting information

Table S1-Plasmids and Strains

Data Set S1

## Acknowledgements

We are grateful to Joaquín Bernal, Pedro Escoll-Guerrero, Rocío López-Igual, José Antonio Escudero, Alexandra Nivina, Celine Loot, Juan Mondotte and Carla Saleh for useful discussions. We thank the technical assistance from: Jean Yves Tivenez for assistance and initial observations in FRAP experiments; Laurence Ma and Christiane Bouchier from the Institut Pasteur Genomics Platform for genomic DNA sequencing; Bertrand Raynal, Sébastien Brulé and Mounira Tijouani for experimental advice on AUC.

This study was supported by the Institut Pasteur, the Centre National de la Recherche Scientifique (UMR3525), the French National Research Agency grants ANR-10-BLAN-131301 (BMC) and ANR-14-CE10-0007 (MAGISBAC), the French Government’s Investissement d’Avenir Program, Laboratoire d’Excellence “Integrative Biology of Emerging Infectious Diseases” (ANR-10-LABX-62-IBEID to DM) and the Agencia Nacional de PromociónCientífica y Tecnológica of Argentina (PICT-2017-0424 to ASB). A.S.-B. was supported by an EMBO long-term fellowship (EMBO-ALTF-1473-2010) and Marie Skłodowska-Curie Actions (FP7-PEOPLE-2011-IIF-BMC). ASB, RS and DJC are Career Members of CONICET. The funders had no role in study design, data collection and analysis, decision to publish, or preparation of the manuscript.

