## Supplementary material for "Macromolecular crowding links ribosomal protein gene dosage to growth rate in *Vibrio cholerae*": Table S1-Plasmids and Strains

**Table S1.** Full list of plasmids, bacteria strains used in this study:

| **Name** | | | **Relevant genotype and/or phenotype** | | **Reference** |
| --- | --- | --- | --- | --- | --- |
| **Plasmids** | | | | | |
| pJBA28 | delivery plasmid for mini-Tn5-Km-PA1/04/03-RBSII-*gfpmut3**-T0-T1, Amp^R^ and Kan^R^ | | | | Andersen et al 1998 |
| pCP20 | pSC101*rep*^TS^ [*flp*] | | | | Cherepanov et al. 1995 |
| pBAD43 | pACYC184 *oriV*, *araBAD*, Spc^R^ | | | | Guzman LM et al. 1995 |
| pRL-CMV | Wild type Renilla Luciferase (RL) gene for expression in mammalian cells. Used for RL amplification. | | | | Promega |
| pGL-2Basic | Wild type Firefly Luciferase (FL) gene for expression in mammalian cells. Used for FL amplification. | | | | Promega |
| pASB12 | pCR-BluntII-Topo(RL gene linked to lox66-Zeo^R^-lox71 cassette) | | | | This study |
| pASB13 | pCR-BluntII-Topo(FL gene linked to lox66 Zeo^R^-lox71 cassette) | | | | This study |
| pASB21 | pCR-BluntII-Topo(*gfpmut3***Not*I fragment-lox66-Zeo^R^-lox71) | | | | Soler Bistué et al 2017 |
| pASB25 | *rpoA* (VC2571) cloned into pBAD43 for its regulated expression. | | | | This study |
| pASB26 | *secY* (VC2576) cloned into pBAD43 for its regulated expression. | | | | This study |
| ***Escherichia coli*** | | | | | |
| DH5α | | | F^-^ *endA1 glnV44 thi-1 recA1 relA1 gyrA96 deoR nupG* Φ80d*lacZ*ΔM15 Δ(*lacZYA-argF*)U169, *hsdR17*(r_K_^-^ m_K_^+^), λ^–^ | |  |
| ***Vibrio cholerae*** | | | | | |
| N16961*ChapRΔlacZ* | | | N16961::mTn*7hapR^+^* Δ*lacZ*. Er^S^, Gn^R^ and Cm^S^. | | Val et al. 2012 |
| Parental -1120 | | | PGB-A192::*attB*’*-lox66-dfrB1-lox71* inserted in the intergenic region between VC1508-VC1509. Er^S^, Gn^R^ and Cm^R^. | | Soler-Bistué et al. 2015 |
| S10Tnp+166 | | | S10 relocated closer to *oriC1* in the intergenic region between VC2739-VC2740. | | Soler-Bistué et al. 2015 |
| S10Tnp-35 | | | S10 relocated next to its original location in the intergenic region between VC2536-VC2537. | | Soler-Bistué et al. 2015 |
| S10Tnp-510 | | | S10 relocated at the middle of the left replichore of chromosome 1 in the intergenic region between VC2075-VC2076. | | Soler-Bistué et al. 2015 |
| S10Tnp-1120 | | | S10 relocated near the *dif* region of chromosome 1 in the intergenic region VC1508-VC1509. Er^S^, Gn^R^ and Cm^R^. | | Soler-Bistué et al. 2015 |
| S10TnpC2+37 | | | S10 relocated near the *oriC2* in the intergenic region between VCA0030-VCA0031. | | Soler-Bistué et al. 2015 |
| S10TnpC2+479 | | | S10 relocated near the *dif* sequence of chromosome 2 in the intergenic region between VCA0543-VCA0544. | | Soler-Bistué et al. 2015 |
| S10Md(-1120;C2+479) | | | Merodiploid bearing *S10-spc-α* copies at the intergenic sequences of VC1508-VC1509 and VCA0543-VCA0544. Er^S^, Gn^R^ and Cm^R^. | | Soler-Bistué et al. 2015 |
| *V.cholerae*::*gfpmut3** | | | NotI fragment from pJBA28 (Andersen et al. (1998)) containing promoter P_A1/04/3_, RBSII and gfpmut3* linked to Zeo^R^ from pASB11 gene was inserted in the intergenic region between VC0696-VC0697. | | Soler-Bistué et al. 2017 |
| Parental-1120::*gfpmut3** | | | *gfpmut3**-ZeoR cassette was inserted in the intergenic region between VC0696-VC0697 in Parental-1120 strain. | | Soler-Bistué et al. 2017 |
| S10Tnp-35::*gfpmut3** | | | *gfpmut3**-ZeoR cassette was inserted in the intergenic region between VC0696-VC0697 in S10Tnp-35 strain. | | This study |
| S10Tnp-510::*gfpmut3** | | | *gfpmut3**-ZeoR cassette was inserted in the intergenic region between VC0696-VC0697 in S10Tnp-510 strain. | | This study |
| S10Tnp-1120::*gfpmut3** | | | *gfpmut3**-ZeoR cassette was inserted in the intergenic region between VC0696-VC0697 in S10Tnp-1120 strain. | | This study |
| S10TnpC2+479::*gfpmut3** | | | *gfpmut3**-ZeoR cassette was inserted in the intergenic region between VC0696-VC0697 in S10TnpC2+479 strain. | | This study |
| Parental-1120::*RL* | | | *RL*-ZeoR cassette from pASB12 was inserted in the intergenic region between VC0696-VC0697 in Parental-1120 strain. | | This study |
| S10Tnp-35::*RL* | | | *RL*-ZeoR cassette from pASB12 was inserted in the intergenic region between VC0696-VC0697 in S10Tnp-35 strain. | | This study |
| S10Tnp-510::*RL* | | | *RL*-ZeoR cassette from pASB12 was inserted in the intergenic region between VC0696-VC0697 in S10Tnp-510 strain. | | This study |
| S10Tnp-1120::*RL* | | | *RL*-ZeoR cassette from pASB12 was inserted in the intergenic region between VC0696-VC0697 in S10Tnp-1120 strain. | | This study |
| S10TnpC2+479::*RL* | | | *RL*-ZeoR cassette from pASB12 was inserted in the intergenic region between VC0696-VC0697 in S10TnpC2+479 strain. | | This study |
| Parental -1120  Δ(*aph,cat*) | | | Parental-1120. Kanamycin and chloramphenicol resistance cassettes were deleted using a flipase expressing plasmid. Er^S^, Gn^R^ and Cm^S^. | | Soler-Bistué et al 2015 |
| S10Tnp-1120 Δ(*aph,cat*) | | | S10 relocated near the *dif* region of chromosome 1. Derived from Parental -1120 Δ(*aph,cat*). Er^S^, Gn^R^ and Cm^S^. | | Soler-Bistué et al 2015 |
| Parental-1120 (pBAD43) | | | Parental strain bearing empty vector pBAD43. | | This study |
| Parental-1120 (pASB25) | | | Parental strain bearing plasmid for *rpoA* overexpression. | | This study |
| Parental-1120 (pASB26) | | | Parental strain bearing plasmid for *secY* overexpression. | | This study |
| S10Tnp-1120 (pBAD43) | | | Strain whose S10 was relocated close to *ter1* bearing empty vector pBAD43. | | This study |
| S10Tnp-1120 (pASB25) | | | Strain whose S10 was relocated close to *ter1* bearing plasmid for *rpoA* overexpression. | | This study |
| S10Tnp-1120 (pASB26) | | | Strain whose S10 was relocated close to *ter1* bearing plasmid for *secY* overexpression. | | This study |
| S10TnpC2+479 (pBAD43) | | | Strain whose S10 was relocated close to *ter2* bearing empty vector pBAD43. | | This study |
| S10TnpC2+479 (pASB25) | | | Strain whose S10 was relocated close to *ter2* bearing plasmid for *rpoA* overexpression. | | This study |
| S10TnpC2+479 (pASB26) | | | Strain whose S10 was relocated close to *ter2* bearing plasmid for *secY* overexpression. | | This study |
| Parental-1120::*gfpmut3** Δ*crts* | | | The Chr2 replication triggering site (*crt*S) a 150 bp Chr1 sequence (coordinates 817950-818100) was replaced by a rifampicin resistance cassette (*arr2*) as in Val et al (2016) Parental-1120::*gfpmut3** | | This study |
| S10Tnp-1120::*gfpmut3** Δ*crts* | | | The *crt*S was replaced by a rifampicin resistance cassette (*arr2*) as in Val et al (2016) in S10Tnp-1120::*gfpmut3** movant. | | This study |
| S10TnpC2+479::*gfpmut3** Δ*crts* | | | The *crt*S was replaced by a rifampicin resistance cassette (*arr2*) as in Val et al (2016) in S10TnpC2+479::*gfpmut3** movant. | | This study |
