## Supplementary material for "Macromolecular crowding links ribosomal protein gene dosage to growth rate in *Vibrio cholerae*": Table S2

**Table S2:** Exponential fit of fluorescence (GFP production) as a function of OD_600nm._  Data was adjusted to the equation Y=Y_0_*exp(k*X).

|  | **Parental** | **S10Tnp-35** | **S10Tnp-510** | **S10Tnp-1120** | **S10TnpC2+479** | ***gfpmut3^-^*** |
| --- | --- | --- | --- | --- | --- | --- |
| **Y_0_** | 2493 | 2507 | 2482 | 2474 | 2475 | 2808 |
| **k** | 2.080 | 2.144 | 2.277 | 2.299 | 2.029 | 0.7047 |
| **R^2^** | 0.9985 | 0.9964 | 0.9968 | 0.9984 | 0.9970 | 0.9844 |
