## Supplementary material for "Macromolecular crowding links ribosomal protein gene dosage to growth rate in *Vibrio cholerae*": Table S4

**Table Salt: Transcriptionally altered genes are shared between movants and regulated in the same manner.** The proportion of altered genes that is also found to be regulated in either of the other two movants is shown in the first column (grey). The percentage is shown in parentheses. The two entry table shows that these altered genes were transcriptionally altered in the same way between the movants. We calculated the Pearson Correlation Test for their Log_2_(FC). The value of the test is shown in green while the corresponding p-value is displayed in orange.

| **S10Tnp** | **All** | **-510** | **-1120** | **C2+479** |
| --- | --- | --- | --- | --- |
| **-510** | 104/111 (93.7%) |  | 0.927 | 0.992 |
| **-1120** | 457/662(69%) | 10^-24^ |  | 0.946 |
| **C2+479** | 501/742 (67.4%) | 10^-30^ | 10^-33^ |  |
