## Supplementary material for "Macromolecular crowding links ribosomal protein gene dosage to growth rate in *Vibrio cholerae*": Table S5

**Table S5: Altered functions upon S10 relocation.** Genes within *V. cholerae* genome were classified in functional categories using eggNOG database. The table shows the number of genes whose expression is altered in selected functional categories for each movant train and the total genes in the chromosome belonging to each category. The number in parenthesis represents the % with respect the total number of genes. The functions with no genes or with no alterations are not displayed and can be found in Data Set 1. The total number in the last row also includes functions not displayed.

|  | **-510** | | | **-1120** | | | **C2+479** | | | **Total** |
| --- | --- | --- | --- | --- | --- | --- | --- | --- | --- | --- |
|  | **down** | **Up** | **Total** | **down** | **Up** | **Total** | **down** | **Up** | **Total** |  |
| J | 2 | 1 | 3  (2.7) | 10 | 10 | 20 (3.1) | 15 | 11 | 26 (3.6) | **172 (5.06)** |
| V | 0 | 0 | 0 | 5 | 7 | 12  (1.86) | 5 | 8 | 13 (1.79) | **48 (1.41)** |
| U | 1 | 3 | 4  (3.6) | 8 | 9 | 17 | 6 | 9 | 15 | **70 (2.06)** |
| O | 1 | 0 | 1  (0.9) | 14 | 32 | 46 (7.14) | 14 | 26 | 40  (5.51) | **121 (3.56)** |
| E | 16 | 9 | 25 (22.5) | 49 | 24 | 73  (11.3) | 40 | 29 | 69  (9.5) | **236 (6.95)** |
| P | 1 | 26 | 27 (24.3) | 18 | 12 | 30  (4.65) | 15 | 35 | 50 (6.9) | **201 (5.9)** |
| **Total** | **49** | **62** | **111** | **296** | **348** | **644** | **296** | **429** | **725** | **3395** |
