## Supplementary Methods for "Macromolecular crowding links ribosomal protein gene dosage to growth rate in *Vibrio cholerae*"

**Supplementary Text**

**Extended methods:**

**Culture conditions:** For fast-growing conditions, bacterial cultures were done in Lysogeny Broth (LB) at 37ºC with maximum agitation. Automated growth curves were performed in 96-well plates avoiding the use of external rows and columns. For harvesting cells in fast growing conditions, ~30 µL of an ON culture was used to inoculate pre-warmed 250 mL Erlenmeyer Flasks with 70 mL of LB and agitation was set to 250 rpm (as in Figs. 2b, 2c, 2e, 3, 4, 6c and 6d). For selection the following antibiotic concentrations were used: chloramphenicol (3 μg/mL), kanamycin (25 μg/mL), spectinomycin (100 μg/mL), carbenicillin (50 μg/mL) and zeocin (25 μg/mL). NaCl and sucrose were added at the indicated concentrations. Strains and plasmids used are listed in Table S1. For strains expressing *secY* and *rpoA* from pBAD43 cells were cultured in LB (for leak expression), in LB supplemented with 1% glucose (expression repression) or LB with 0.2% arabinose (induction).

**RNA preparation and sequencing for transcriptomic studies:** 20 mL of and early exponential phase culture were recovered by centrifugation at 4,500 rpm for 10 minutes at 4ºC. Then, RNA was extracted using Trizol (Thermo Fisher Scientific). Residual DNA was removed with TURBO DNAse (Ambion). RNA quality (total, depleted, purified) was checked on the Bioanalyzer 2100 (Agilent). Samples were cheked for RNA Integrity Number >8. The rRNA was depleted using the MicrobExpress kit (Ambion) and libraries were built using TruSeq Stranded RNA LT Sample Prep Kit (Illumina) and checked for concentration and quality on Bioanalyzer and QuBit (Invitrogen). Sequencing of multiplexed libraries was performed on a HiSeq 2500 (Illumina). Then, in-house quality control process was applied to reads that passed the Illumina quality filters (raw reads). The sequences of the Illumina adapters and primers used during the library construction were removed from the whole reads. Low-quality nucleotides were removed from both ends.

**Marker frequency analysis (MFA) and slope calculation:**

Genomic DNA extracted from early exponential phase (OD_450 nm_ ~0.15) was used for library preparation using a PCR-free protocol. Libraries were sequenced on an Illumina MiSeq sequencer using 100- to 150-base-length paired-end reads for 100× genome coverage. The resulting trimmed FastQ files were analyzed using R2R script to obtain the frequency of each locus along the genome, removing repeated sequences [1-3]. Values were normalized against from a stationary phase of a parental strain control for noise reduction. Then, the Log_2_ frequencies every 1,000-bp window were then plotted as a function of their relative position on chromosome 1 in *ter1-ori1-ter1* order (Figure 3b). Slopes were obtained from linear regression of plots of Log_2_ frequencies along replichore length from ter1 to ori1 (R^2^>0.95). The slopes represent the log2 frequency change per Kbp (Figures 3c, S5 and Table S3).

The frequency of ori1 and ter1 were quantified by averaging 50 frequency data points corresponding to ori1 and ter1 zones. The S10 frequency was calculated by averaging panels corresponding to VC2569 and VC2599, respectively. These values were used to calculate S10 dosage with high precision by calculating the S10/ter1 ratio (Figure S5 and Table S3).

**COG CATEGORIES:**

INFORMATION STORAGE AND PROCESSING

[J] Translation, ribosomal structure and biogenesis

[A] RNA processing and modification

[K] Transcription

[L] Replication, recombination and repair

[B] Chromatin structure and dynamics

CELLULAR PROCESSES AND SIGNALING

[D] Cell cycle control, cell division, chromosome partitioning

[Y] Nuclear structure

[V] Defense mechanisms

[T] Signal transduction mechanisms

[M] Cell wall/membrane/envelope biogenesis

[N] Cell motility

[Z] Cytoskeleton

[W] Extracellular structures

[U] Intracellular trafficking, secretion, and vesicular transport

[O] Posttranslational modification, protein turnover, chaperones

METABOLISM

[C] Energy production and conversion

[G] Carbohydrate transport and metabolism

[E] Amino acid transport and metabolism

[F] Nucleotide transport and metabolism

[H] Coenzyme transport and metabolism

[I] Lipid transport and metabolism

[P] Inorganic ion transport and metabolism

[Q] Secondary metabolites biosynthesis, transport and catabolism

POORLY CHARACTERIZED

[R] General function prediction only

[S] Function unknown

**RNA-Seq Statistical analysis:** Count data were analyzed using R version 3.1.2 [4] and the Bioconductor package DESeq2 version 1.6.1 [5]. Data were normalized with DESeq2 and the “shorth” parameter. The dispersion estimation and statistical test for differential expression were performed with default parameters (including outlier detection and independent filtering). The generalized linear model was set with strain (Parental, S10Tnp-1120, S10Tnp-35, S10Tnp-510 and S10TnpC2+479 levels) as main effect. Since samples were prepared 4 times independently, the date of sample preparation was also included into the model as a blocking factor to catch more variability and increase the statistical power. Raw p-values were adjusted for multiple testing according to the Benjamini and Hochsberg (BH) procedure [6] and genes with an adjusted p-value lower than 0.05 were considered differentially expressed.

**Analysis of FRAP images:**

For long-term experiments, (GFP re-synthesis) detection images were taken every 2 seconds for at least 5 minutes using 200-500ms of acquisition time. Image analysis was done using ImageJ following photobleached and non-bleached cells in time. The average signal of not-photobleached cells was subtracted to the signal of bleached cells to take into account the decay produced by cell imaging.

For measurement of GFP diffusion within bacteria, we used the exposure times 20ms at maximum acquisition frame rate. Then these movies were analyzed using a specific Jython script developed during the Image Processing School Pilsen 2009 and updated to modern Fiji as described in [7]. For every cell analyzed cell length, the photobleached area and the total cell area were determined. Also a control area was measured. Then the Jython script was executed. For data analysis we only kept cells shorter than 6 µm. We only registered half-time values when the function fitted with R^2^ > 0.8.

To discard that half-time recovery of fluorescence (τ) differences observed were not due to an unintentionally biased analysis we measured and analyzed cell length, cell area, control area, photobleached area and the mobile fraction. As explained in the text, τ was longer in the parental strain. The length and the area of the cell were significantly lower in C2+479. The photo bleached area was not significantly different among strains although there is a trend for parental strain to present a larger one.

|  | Parental | C2+479 | -1120 |
| --- | --- | --- | --- |
| - (ms) | (120.4-158.9) | (97.39-117.52) | (88.31-106.3) |
| cell lenght (µm) | (4.87-5.25) | (4,35-4,72) | (4,85-5,2) |
| Photobleached area (µm^2^) | (1.74-1.95) | (1,51-1,80) | (1,56-1,75) |
| Control Area (µm^2^) | (1.64-1.9) | (1,46-1,73) | (1,67-1,87) |
| Cell Area (µm^2^) | (4.32-4.73) | (3,93-4,32) | (4,07-4,43) |
| Mobile faction (%) | (22.37-25.98) | (21,25-25,21) | (22,33-25,16) |

We next performed spearman correlation analyses between these variables among these strains. Interestingly, cell length correlated with τ in all 3 strains (r= 0.487, r=0.49 and r=0.399 respectively with p<0.0001 in all cases). The photo bleached area presented small r value (~0.2) that was not statistically significant in the parental strain. Therefore, we consider the influence of the bleached area in the obtained τ very mild or inexistent. Finally, we found that the employed control area did not influence the outcome of the experiments since there was not a correlation between τ and this parameter.

[4] R Core Team, R: A Language and Environment for Statistical Computing, R Foundation for Statistical Computing, 2014.

[5] Love, Huber and Anders, Moderated estimation of fold change and dispersion for RNA-Seq data with DESeq2, Genome Biology, 2014.

[6] Benjamini and Hochberg, Controlling the False Discovery Rate: A Practical and Powerful Approach to Multiple Testing, Journal of the Royal Statistical Society, 1995.

[7] https://imagej.net/Analyze_FRAP_movies_with_a_Jython_script
