## Supplementary material for "Macromolecular crowding links ribosomal protein gene dosage to growth rate in *Vibrio cholerae*": Figure S1

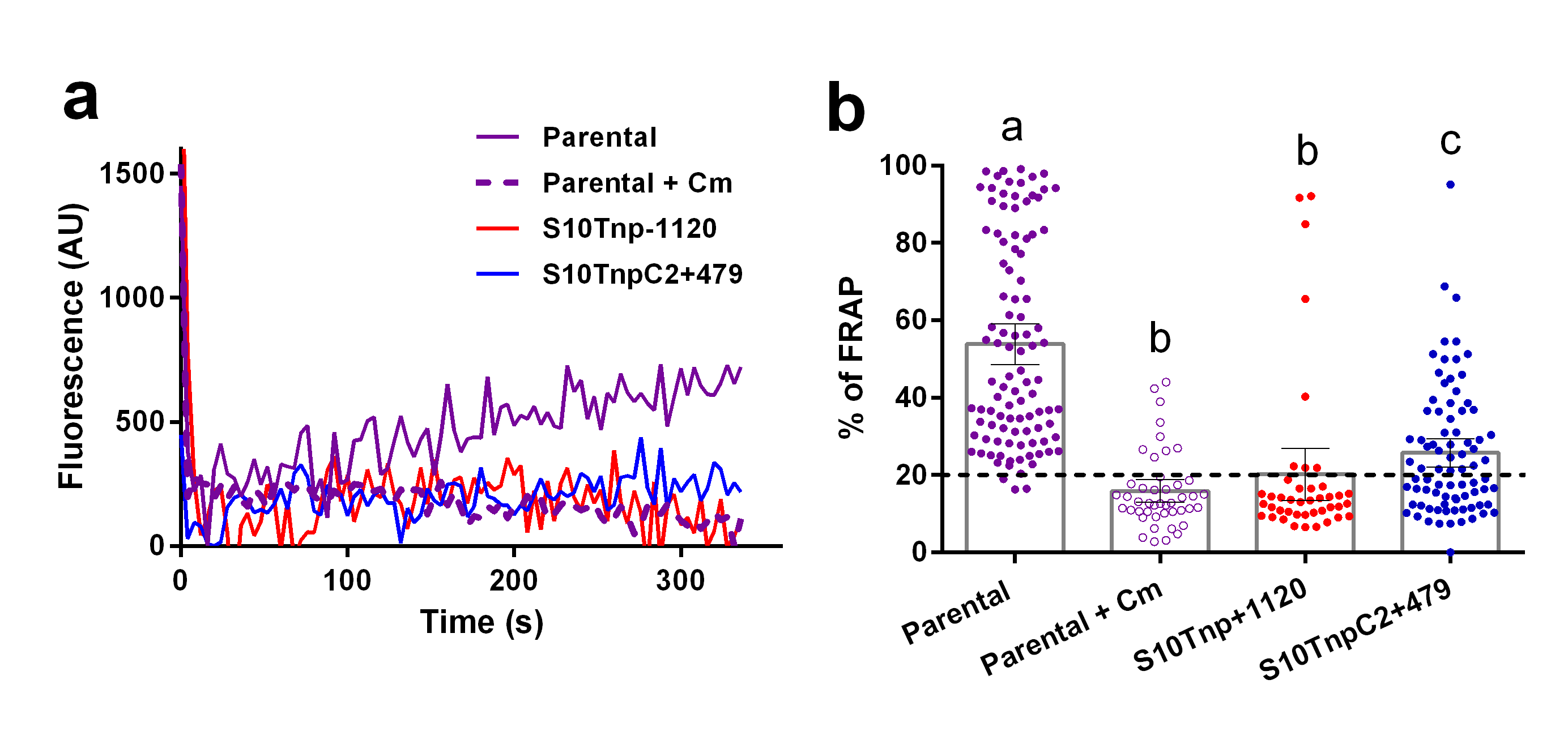

**Figure S1: The most affected movants, display lower GFP production than the Parental strain at the single cell level.** FRAP experiments were performed in LB at 37°C taking a photo every 2 seconds for at least 5 minutes using the Parental-1120 strain (Parental, violet), the S10Tnp-1120 (red) and S10TnpC2+479 (blue) movants. The parental was also tested in presence of chloramphenicol at MIC (+ Cm). **a)** A representative plot showing the recovery of fluorescence over time in individual cells. **b)** The percentage of FRAP at the endpoint of the experiment is shown for all cells tested. Mean with 95% CI is shown. Statistical significance was analyzed by Kruskal-Wallis test (p<0.0001). Then Dunn multiple comparison test was made for mean rank obtained for each strain. Letters denote groups being statistically different.
