## Supplementary material for "Macromolecular crowding links ribosomal protein gene dosage to growth rate in *Vibrio cholerae*": Figure S2

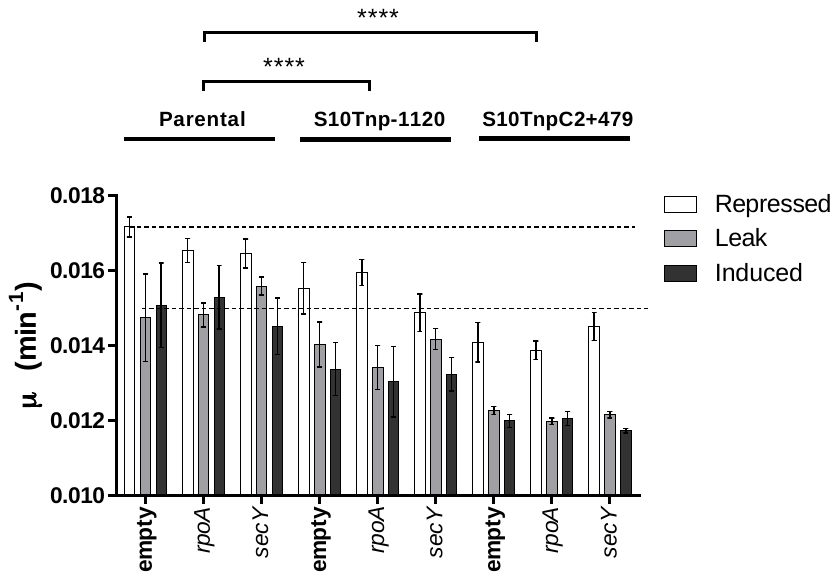

**Figure S2: *rpoA* and *secY* overexpression does not rescue growth rate impairment due to S10 relocation.** Effect of empty vector, *rpoA* or *secY* expression was quantified by averaging the slope (μ) obtained using 4 biological replicates for each strain in different induction conditions. Results are expressed as the mean μ ± 95% CI. Statistical significance was analyzed using a two-way ANOVA two tailed test and Tukey test for multiple comparisons (p<0.0001). Independently of culture conditions differences are not statistically significant between strains harboring the empty vector, pASB25 or pASB26. Expression was repressed by supplementing culture media with 1% glucose. Induction was achieved adding L-arabinose up to 0.2%.
