## Supplementary material for "Macromolecular crowding links ribosomal protein gene dosage to growth rate in *Vibrio cholerae*": Figure S4

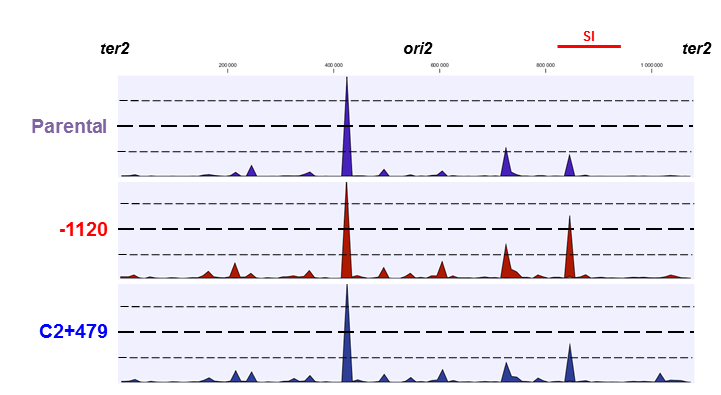

**Figure S4: RNA Coverage of chromosome 2 (Chr2) on selected strains.** RNA prepared in fast-growing conditions was subjected to deep sequencing. Reads were mapped along the Chr2 of *V. cholerae*. The graphs show Normalized Expression Values along both replichores of the replicon in *ter2-ori2-ter2* order of the parental and the most affected strains. Each graph represents the coverage along Chr2 length of Parental (purple), S10Tnp-1120 (red) and S10TnpC2+479 (blue). The superintegron is highlighted in red (SI). Interestingly, SI region is overexpressed in the S10Tnp-1120 movant. Scale from the first base is shown above the graphs.
