## Supplementary material for "Macromolecular crowding links ribosomal protein gene dosage to growth rate in *Vibrio cholerae*": Figure S5

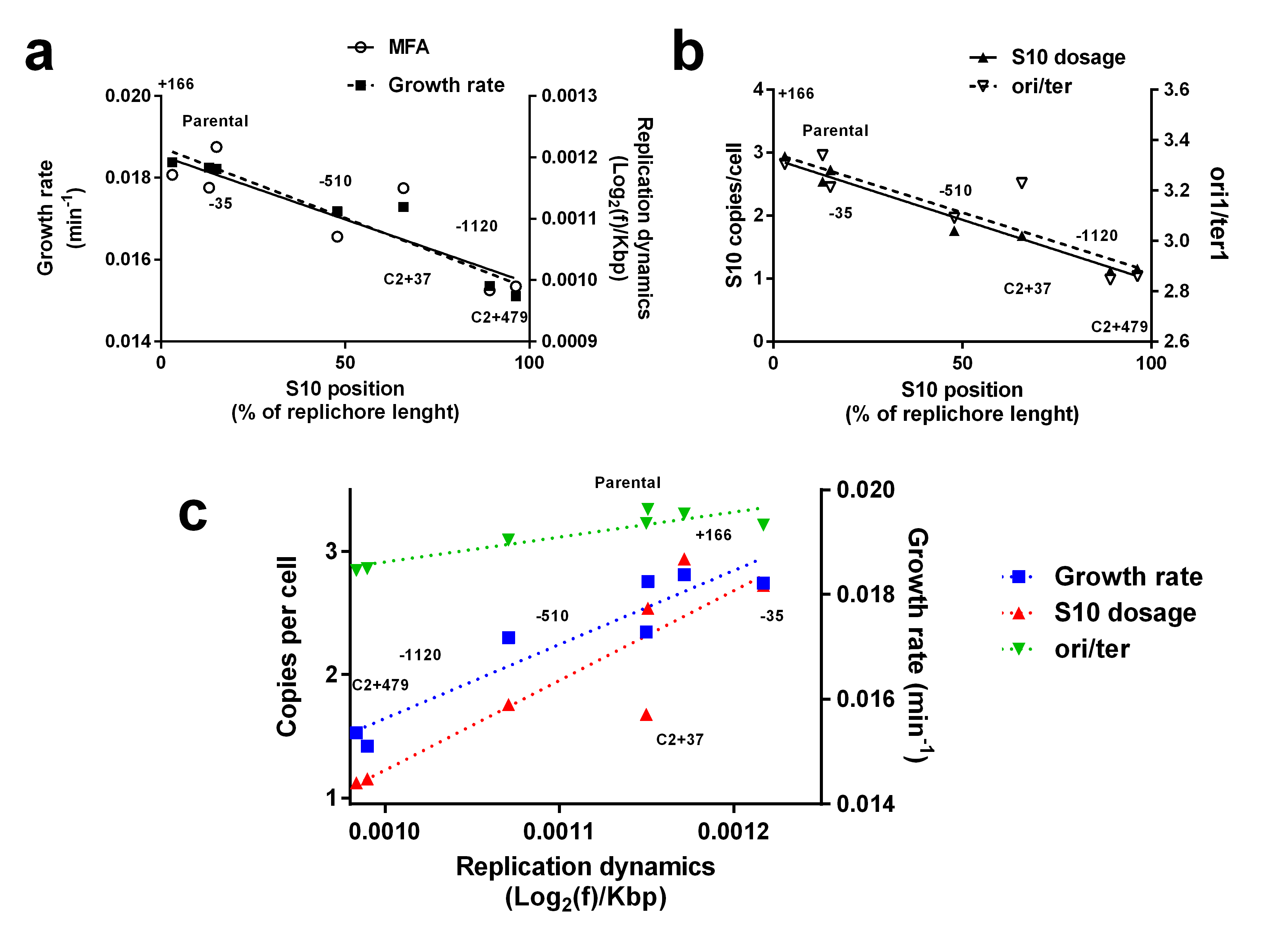

**Figure S5: Replication dynamics closely correlates S10 location, dosage, *ori1* firing and growth rate.**  **a)** The slopes obtained from the MFA analyses (white circles, right axis) and the growth rate (black squares, left axis) of each strain were plotted as a function of the S10 genomic location. **b)** S10 dosage (black triangles, left axis) and ori1/ter1 (white triangles, right axis) ratio from MFA analyses for each strain were graphed as a function of the S10 positioning. **c)** S10 dosage (red), ori1/ter1 ratio (green) and growth rate (blue) are plotted as a function the slope obtained for each strain in MFA analyses. Linear regression for each variable is shown in dotted lines. The data used for each graphic can be found in Table S3. The obtained correlations and their statistical significance are described in the main text of the article.
