## Supplementary material for "Macromolecular crowding links ribosomal protein gene dosage to growth rate in *Vibrio cholerae*": Figure S6

**Figure S6: S10 relocation produces homogeneously distributed global changes in *V. cholerae* gene expression.** Circos plot of genome-wide expression data from strains S10Tnp-35 (Turquoise), S10Tnp-510 (green), S10Tnp-1120 (red) and S10TnpC2+479 (blue). Upper case represents Chr1 while lower case is Chr2 in ter-ori-ter disposition. The origin of replication of each chromosome is represented as *oriC1* and *oriC2* respectively. From inside to outside: Sense and antisense *V. cholerae* genes are depicted as dark orange and orange boxes, respectively. Blue bars represent RNA-seq read counts per gene (scale 1-200,000). Fold-change expression relative to the parental strain is indicated as a green or red solid line indicating fold-expression differences higher than 1.2 or lower than 0.8, respetcively. Dark red dots indicate –log (*p-*value) of the differential expression analysis. Notably, the abundance of significantly altered genes (red dots) from left to right.

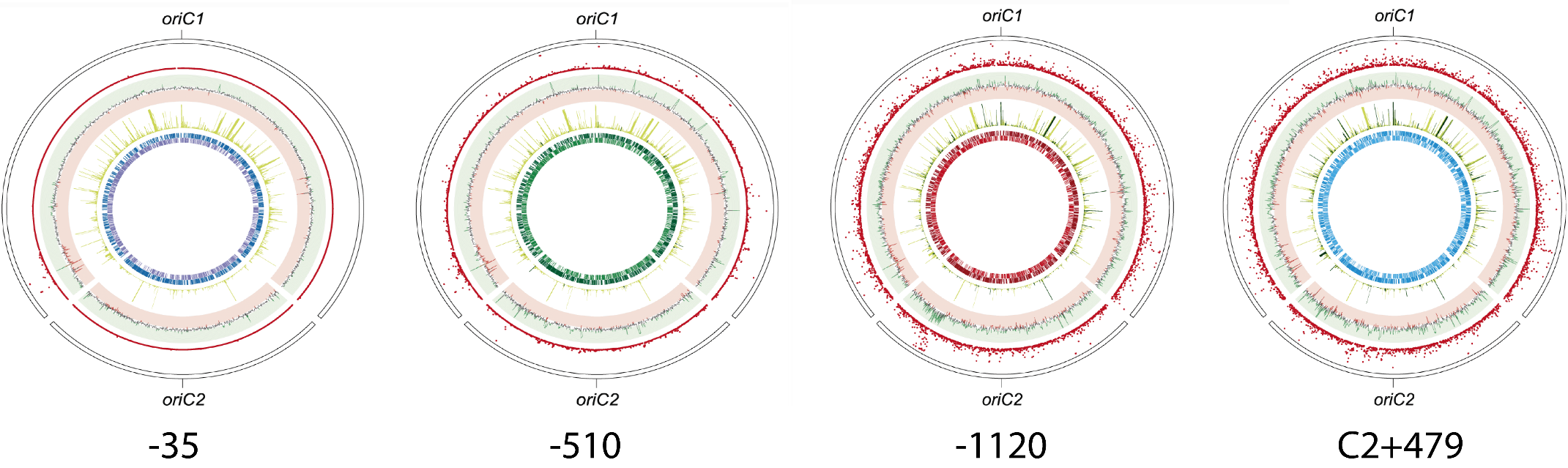
