## Supplementary material for "Macromolecular crowding links ribosomal protein gene dosage to growth rate in *Vibrio cholerae*": Figure S7

**Figure S7:** Manhattan Pot showing statistically altered functions across the movant strain set. The abscissa correspond to specific COG within the S10Tnp-510 (green), S10Tnp-1120 (red) and S10TnpC2+479(blue). S10Tnp-35 is not included since very few genes are differentially expressed displaying no altered functions. The purple line indicates statistical significance fixing α in 0.05.

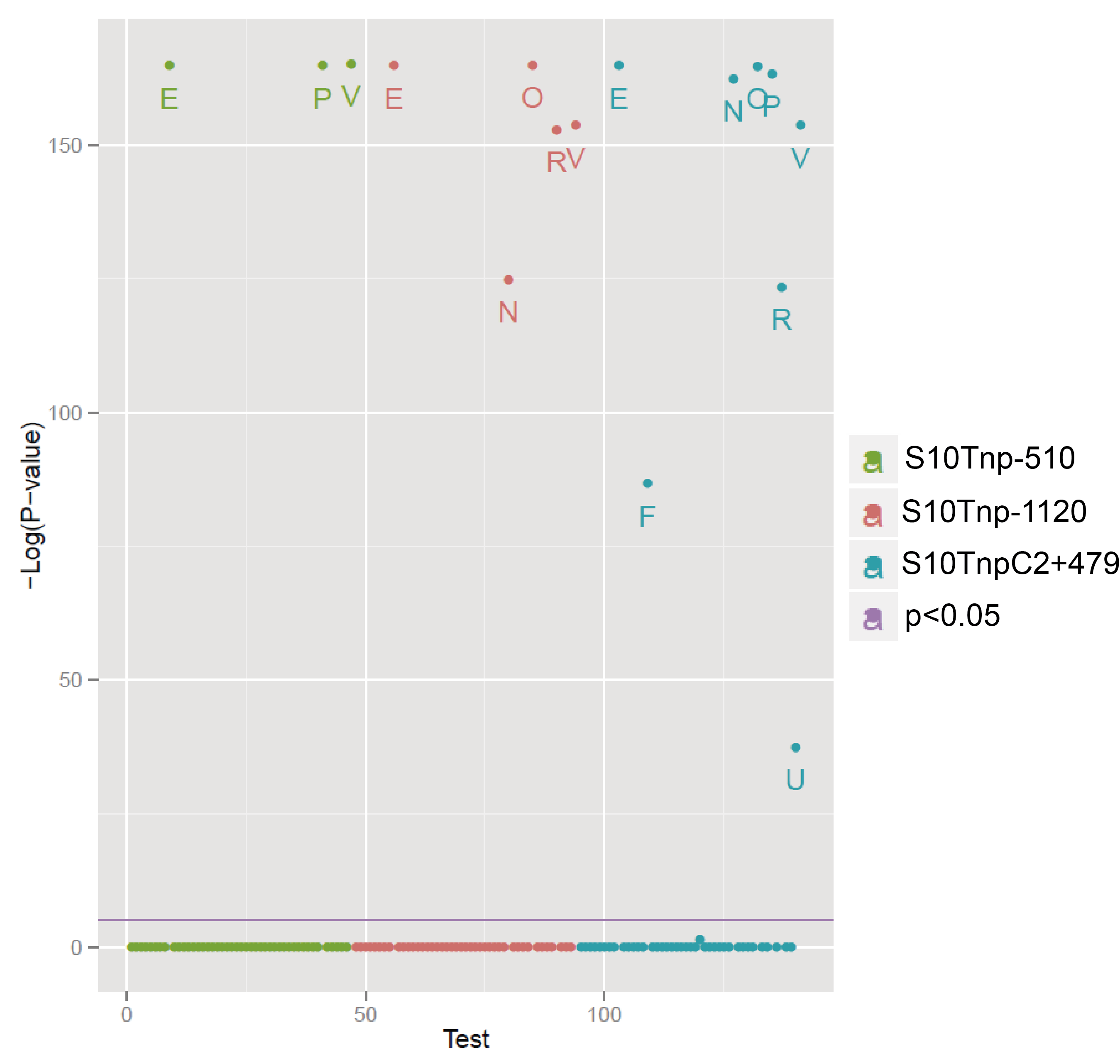
