## Supplementary material for "Macromolecular crowding links ribosomal protein gene dosage to growth rate in *Vibrio cholerae*": Figure S8

**Figure S7:** The growth rate of the parental strain, S10Tnp-1120 and S10TnpC2+479 was measured using automated growth curves at different NaCl concentrations in rich medium. The mean µ value with SEM of 5 independent experiments is shown. Statistical significance was analyzed by one-way ANOVA two-tailed test. Then Holm-Sidak test was done to compare the means values obtained for each strain. Letters denote groups being statistically different within strains. Differences between strains within each NaCl concentration are denoted as follows: *, p<0.05; **, p<0.01; ***, p<0.001 and n.s. stands for non-significant.

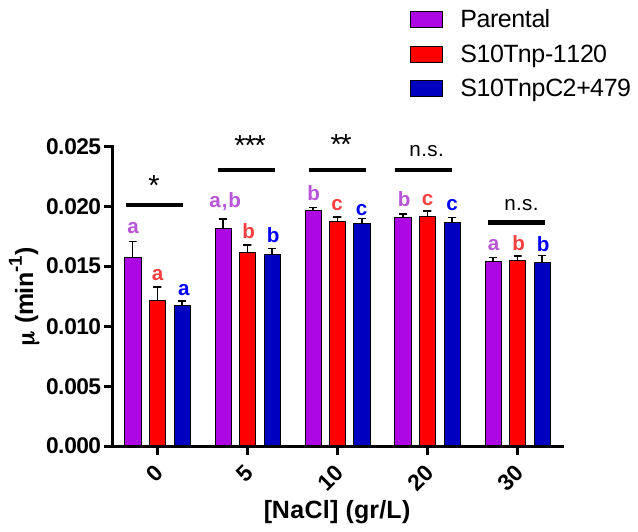
