## Supplementary material for "Macromolecular crowding links ribosomal protein gene dosage to growth rate in *Vibrio cholerae*": Figure S3

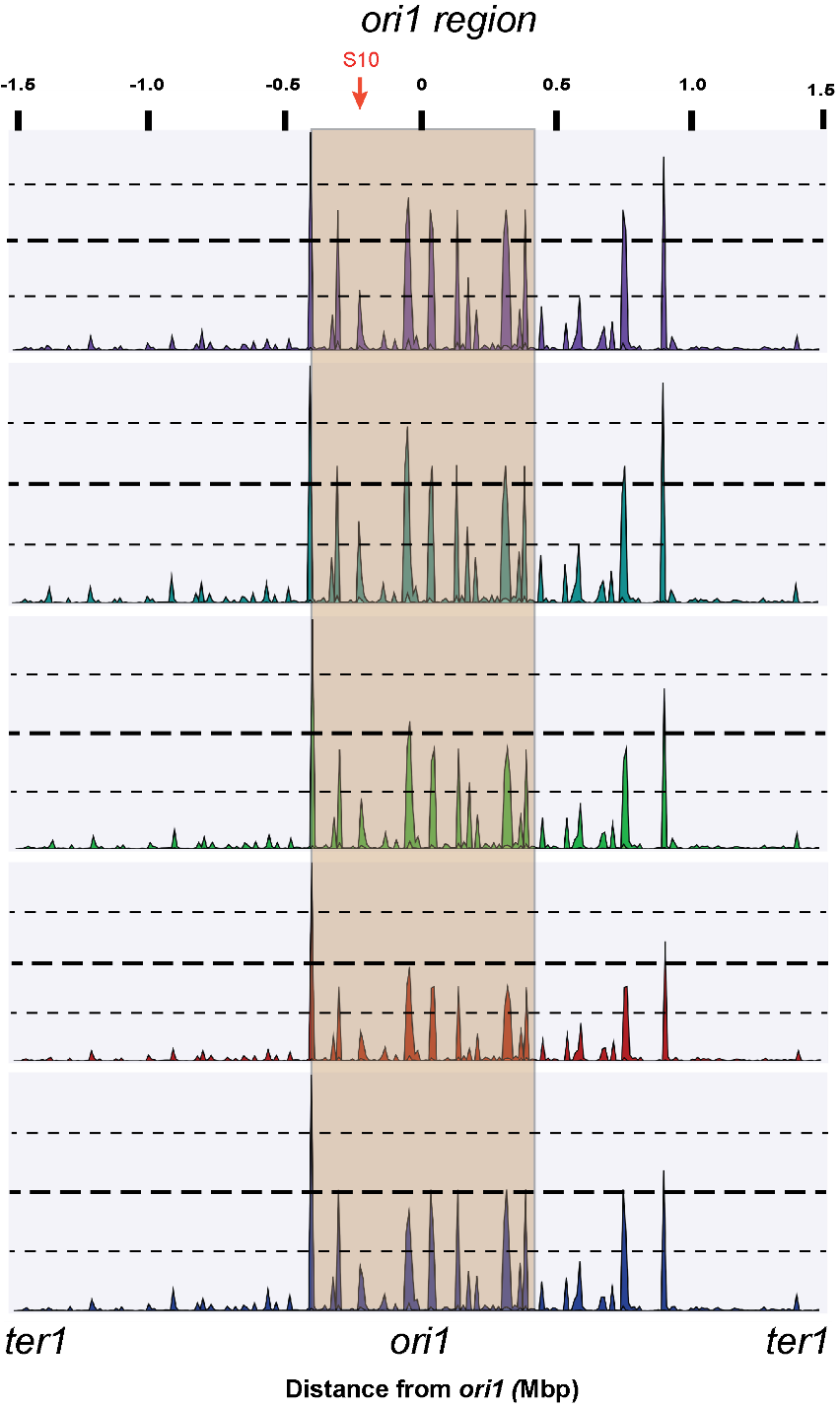

**Figure S3: RNA Coverage of Chromosome 1 (Chr1) on the full movant strain set.** RNA prepared in exponential phase was Deep-Sequenced as described in Materials and Methods. Reads were mapped along the Chr1 of *Vibrio cholerae* and normalized against the full sequence volume. The graphs show the coverage as Normalized Expression Values (dotted lines indicate 75, 50 and 25e10^3^ NEV) along both replichores of the replicon in *ter1-ori1-ter1* order. Each graph represents one strain: Parental (purple); S10Tnp-35 (cyan); S10Tnp-510 (green); S10Tnp-1120 (red); S10TnpC2+479 (blue). The 400 Kbp flanking ori1 are highlighted in orange. A red arrow indicates the peak corresponding to the S10 locus. The coverage of the *ori1* region and the size of the S10 peak lowers with increasing S10-*ori1* distance (see text and Fig. 3a).
